# How informative are structural alphabets for representing protein folds?

**DOI:** 10.64898/2026.01.16.699908

**Authors:** Ferras El-Hendi, Ali Al-Fatlawi, Ballal Hossen, Tom Kasper, Alexandre Mestiashvili, Michael Schroeder

**Author notes:** These authors contributed equally to this work (Ferras El-Hendi, Ali Al-Fatlawi). Department of Surgery, Friedrich-Alexander-Universität Erlangen-Nürnberg and Universitätsklinikum Erlangen, Erlangen, Germany (Ali Al-Fatlawi).

## Abstract

Amino acid sequences are the primary basis for protein comparison, but sequence similarity is often difficult to detect between distantly related proteins, even whe folds remain conserved. Therefore, comparing three-dimensional (3D) protein stru can reveal relationships that are not apparent from sequence alone. Full structural alignment uses 3D coordinates, whereas structural alphabets represent protein structures as sequences of discrete states. These alphabets differ in granularity: represents helices, strands, and loops using three states; Q8 uses the eight secondary-structure states defined by DSSP; and the 20-state 3Di alphabet, dev for Foldseek, describes tertiary residue-residue interactions. Here, we quantified fold-level information retained by Q3, Q8, and 3Di across three CATH and SCO fold-recognition benchmarks, using TM-scores as the reference measure of full-str similarity. The results reveal a Pareto-like trade-off: despite reducing protein str to a linear pattern of helices, strands, and loops and discarding explicit information about spatial distances, residue orientations, and tertiary contacts, Q3 preserves approximately 90-96% of the global fold-recognition signal available from full-st comparison and is on par with the richer 20-state 3Di alphabet. Although Q3 lac fine-grained structural details needed to capture the remaining signal, it provide simple, interpretable, and information-rich representation of protein-fold similarit central purpose of this study is therefore to quantify the fold-level information re by a three-state secondary-structure representation, rather than to introduce Q3 alternative to full structural comparison or optimized search methods.

## 1 Introduction

### Sequence and structure in remote homology detection

Homologous proteins share a common evolutionary origin and may retain similarities in sequence, structure, and function. Amino-acid sequence comparison is the primary basis for detecting such relationships, using methods ranging from pairwise searches such as BLAST [2] to profile-based approaches such as HHblits and HHsearch [3, 4]. However, sequence similarity is often difficult to detect between distantly related proteins, even when their folds remain conserved. Because three-dimensional (3D) protein structure is generally more conserved than sequence [5–7], structural comparison can reveal relationships that are not apparent from sequence alone.

Coordinate-based structural alignment compares proteins using 3D coordinates and provides a detailed reference for fold-level similarity; in this study, this similarity is summarized using TM-scores [8]. The rapid expansion of predicted-structure resources, including the AlphaFold Protein Structure Database [9] and the ESM Metagenomic Atlas [10], makes it important to understand how much structural information is preserved when 3D structures are compressed. Structural alphabets provide such a compression by representing protein structures as sequences of discrete states that can be compared using sequence-alignment methods.

### Structural alphabets and fold-information retention

Structural alphabets differ in granularity. Q3 represents proteins using three secondary-structure states: helices (H), strands (S), and loops (L). Q8 distinguishes eight states: *α*-helices (H), G-helices (G), *π*-helices (I), *β*-bridges (B), extended *β*-strands (E), bends (S), turns (T), and coils or unassigned residues (C or blank). The 20-state 3Di alphabet, developed for Foldseek, instead uses 20 letters to encode tertiary residue-residue interactions [11]. Previous studies have shown that secondary structure carries useful information for fold recognition, structural alignment, motif identification, and profile-based homology detection [4, 12–17]. However, it remains unclear how much fold-recognition signal is retained when the full 3D structure is reduced to Q3 and whether richer alphabets provide commensurate gains.

The bacterial RedBeta and human Rad52 single-strand annealing proteins provide an illustrative case of remote homology (see Fig. 1). Their structures share a three-stranded beta sheet followed by a packed helix [1], whereas the HHblits alignment covers only a short sequence region. A Q3 alignment makes the shared secondary-structure pattern explicit in a compact and interpretable form. This example motivates, but does not establish, the general information content of Q3.

**Fig 1.**
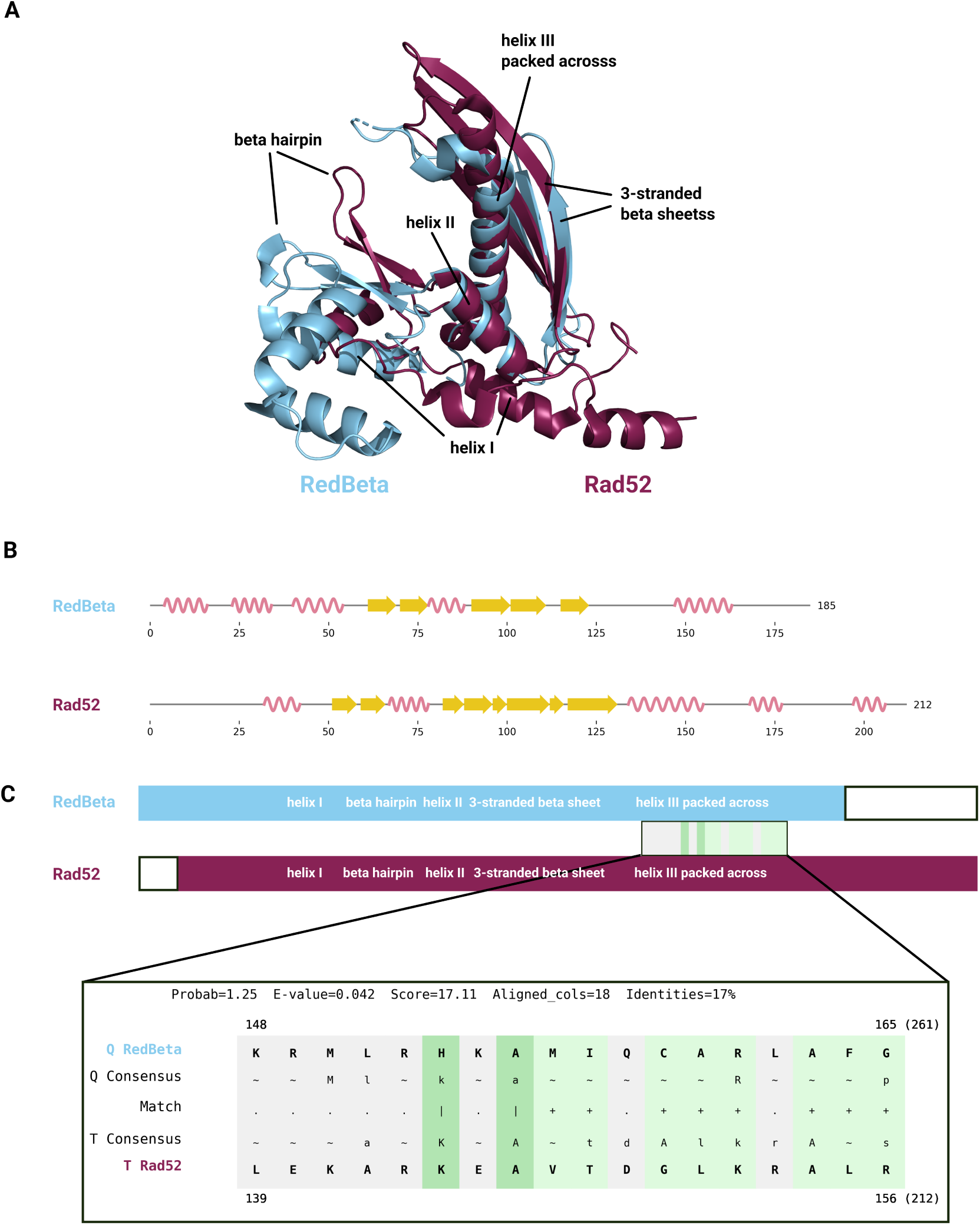
Overview of alignments for Rad52 and RedBeta. **A**: Structural alignment of Rad52 and RedBeta shows clear structural similarity, in particular in those regions, that we can see aligned in the secondary structure alignment. However, due to flexibility, the beta hairpin motif does not get aligned perfectly, nor does the beta sheet due to varying lengths. **B** Alignment of secondary structure. Lengthwise alignment of secondary structure uncovers the shared motifs: a short helix followed by a beta hairpin that connects via a helix to a three-stranded beta sheet and is followed by another helix packed across. This is the shared motif as described in [1]. **C**: HHblits alignment of HMM profiles for Rad52 and RedBeta. Only 18 columns in total around helix III could be aligned (gray-green box) and only 10 with good matching columns (indicated by | and +) and the probability estimate of the pair being homologues is put at 1.25 %. Note that total sequence length (in brackets) is longer than for the secondary structure alignment since not the whole sequence is structurally resolved.

### Study objective and scope

We therefore address a representation-level question: how much global fold-recognition signal is retained when protein structure is reduced from full 3D coordinates to Q3, and how much additional signal is gained by Q8 or 3Di? We evaluate these alphabets on three CATH and SCOPe fold-recognition benchmarks with increasing sequence-identity filtering [18, 19], using TM-scores as the reference measure of full-structure similarity. Each benchmark tests whether two protein domains belong to the same superfamily. Global AUC is the primary measure of representation fidelity, whereas AUROC1 is interpreted separately as a measure of early retrieval. HHblits and Foldseek are included only as contextual end-to-end baselines because their performance also reflects alignment algorithms, scoring schemes, statistical calibration, and database-search procedures. The central contribution is therefore to quantify information retention under extreme structural simplification, not to introduce an alternative to full structural comparison or optimized search methods. We additionally examine the relationship between Q3 and 3Di and present the newt-genome analysis as an illustration of structural-match prioritization rather than validated function annotation.

## 2 Materials and methods

### 2.1 Datasets and baseline methods

CATH version 4.3.0 and SCOPe version 2.07, including the non-redundant datasets CATH S20 and SCOPe40 were used. CATH S20 and SCOPe40 were fully considered; the redundant CATH dataset was filtered to exclude very short (*<*50 residues) and very long (*>*250 residues) domains as well as non-continuous domains.

We obtained the classification information and domain selection criteria from the CATH-Gene3D database version 4.3.0 for our analysis. This release includes data on 500,238 domain classifications. The number of amino acids per domain varies, ranging from a minimum of 6 to a maximum of 1,434, with a mean of 165 (see S4 Note). Based on this data, we defined a representative range of amino acids per domain to be between 50 and 250 amino acids. To further refine our selection, we only considered domains that fell within a single selection range and had the correct residue range associated with a superfamily classification in CATH. Applying these criteria yielded a total of 22,945 domains. We used PyMOL (v2.2.0, Open Source) to trim the domains according to their selection ranges from the corresponding PDB structures, and we saved the resulting domains in .cif format.

For the experiment using predicted Q3 sequences, we used the ProtTrans [20] `ProtT5-XL-U50` model to predict secondary structure using the template provided by the authors on GitHub as reference.

We used BLAST [21] (with default parameters, against each respective dataset), HHblits [3], and TM-Align [22] for basic sequence alignment, advanced sequence alignment, and structure alignment, respectively. For TM-align we used the minimum of both reported scores, i.e. normalised by the longer sequence, to not overestimate TM-align.

For HHblits we first ran all protein domains for each dataset against UniRef30 (2022/02) [23] with E-value cutoff 0.001, max filter 10^8^ and 3 iterations to generate MSAs. Next we used HHmake to turn these MSAs into HMM profiles, and lastly we ran pairwise alignments of the HMM profiles for all domains using HHalign. Finally, we use HHalign’s probability output as scoring measure.

The Foldseek score we used is the bitscore. We used Foldseek with default settings and E-value cutoff 10^300^, to maximize retrieval and enable Foldseek to produce a score for each pair, as does every other method, for each respective dataset. The Foldseek command we ran was:

~~~
foldseek easy-search query.pdb target.pdb query_vs_target_type2.tsv tmp_type2
--alignment-type 2 --exhaustive-search 1 -e 1e300
~~~

AUCs were computed with scikit-learn (roc auc score) over the normalised Levenshtein and normalised Smith-Waterman scores, respectively (see below).

AUROC1 was computed as defined in [11]. AUROC1 plots show cumulative distribution of sensitivity up to the first false positive (FP). True positives (TPs) are matches of the same superfamily, false positives (FPs) those of different fold.

### 2.2 Alphabets and Similarity

Amino acid sequences were translated into the 3-, 8-, and 20-letter alphabets using PyMol (v 2.2.0 Open-Source) [24], DSSP [25] and Mini3Di [26], respectively.

For PyMol, only C*α* atoms were considered. Sequences over these alphabets were compared using the Levenshtein Python C extension module (v 0.20.9). Levenshtein distances *Lev*(*a, b*) between sequences *a* and *b* were normalised as follows: *Lev score*(*a, b*) = 1 *− Lev*(*a, b*)*/max*(*len*(*a*)*, len*(*b*)) We further implemented local alignments for secondary structure with the Smith-Waterman algorithm.

This entailed the definition of gap penalties and a substitution matrix. For the gap penalties, we experimented with values mirroring the BLAST settings and found a penalty of -11 for opening a gap and -3 for extending a gap as optimal. The construction of the substitution matrix followed a similar procedure as BLOSUM [27], the calculation of which derived a log odds score from counts of matches and mismatches. We used Pfam-A Seed alignments [28] as basis for these counts. For 19,226 alignments over a total of 1,155,996 sequences for which AlphaFold structures existed, we derived secondary structure. In accordance with the original BLOSUM authors [27], the alignments were then trimmed to remove columns containing gaps, leaving 2,460,186 ungapped columns (for an analysis of matches/mismatches vs. gaps, see also S5 Note).

With the data prepared, we apply the same steps as for BLOSUM, resulting in the final secondary structure substitution matrix (see S1 Note), which penalises mismatches between helix and strand most heavily and rewards matching strands the highest.

### 2.3 Newt genome protein sequences

The newly sequenced Iberian ribbed newt genome (GenBank Accession: GCA 031143425.1), contains 33,854 candidate genes, which match protein-coding regions in other organisms at significant E-values [29]. From these candidates, we randomly selected 99, for which secondary structure could be predicted with ProtTrans and tertiary structure with the AlphaFold3 web server [30]. These 99 proteins were compared against all 496,866 protein in SwissProt [31], for which there was a high-quality (average pLDDT above 70) predicted structure in the AlphaFoldDB in May 2024 [32, 33] and for which secondary structure could be derived from PyMOL.

### 2.4 Sequence comparison for function annotation

Secondary structure sequences were aligned with a dedicated implementation of the Smith-Waterman algorithm [34] for secondary structure with penalties for gap opening set to -11 and for extension to -3 as well as a dedicated secondary structure substitution matrix (HH 2, LL 3, SS 4, HL -7 HS -16, SL -5, where H is helix, S strand, and L loop, see S1 Note). The raw Smith-Waterman score *SW* (*a, b*) for sequences *a* and *b* was normalised as follows: 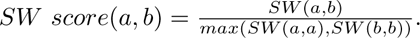 The secondary structure substitution matrix was calculated from high-quality Pfam-A Seed alignments [28] (downloaded 23/04/24), which were translated to secondary structure. Matches and mismatches in these alignments were counted and a log-odds score was calculated following BLOSUM [27]. The affine gap penalties were tuned on a small random sample dataset of 10 proteins from the Newt candidate set, that were different from the 99 proteins we used in the study, and compared against SwissProt.

### 2.5 Function annotation task

Thus, for the 99 newt proteins and for the 496,866 proteins in the screening library, primary, secondary, and tertiary structure were present. Then all 49,189,734 pairs of newt and library proteins were compared with each other using BLAST, PSI-BLAST, the Smith-Waterman algorithm for secondary structure, and US-Align [8].

In a second function annotation study, we selected 20 candidate genes (see S2 Table, S2 Fig) from the newt genome for which the authors could not find protein-coding regions in another organism at a significant E-value [29]. These more challenging candidates were translated, and secondary and tertiary structure predicted as above. Next, their secondary structure was screened against 496,866 proteins in Swiss-Prot. For the best hits, tertiary structure was compared using US-Align [8], as it improves the original TM-align for full proteins.

### 2.6 Posterior Probabilities

We followed the workflow of [35] for calculating posterior probabilities. We used SCOPe40 as the underlying dataset.

### 2.7 Visualization

PyMOL (v 2.5.0) was used to visualise the structures. The Biotite python package (v 0.37.0) has been used to visualise the secondary structure aligned strings [36].

## 3 Results

### 3.1 Fold recognition benchmarks and evaluation measures

To answer how encoding structures as sequences over a multi-state structural alphabet impacts remote homology detection, we consider established fold recognition data sets from the CATH [18] and SCOPe [19] domain classification databases. Given a set of superfamilies with domains from these databases, we compare scores within a superfamily and between superfamilies. The former should be substantially better than the latter.

The difficulty of such a fold recognition task is tightly linked to the amount of sequence redundancy within each superfamily. The less redundant sequences within a superfamily exist, the more difficult it is for a method to infer superfamily membership from a direct sequence comparison. Therefore we employ three datasets: CATH, SCOPe40, and CATH S20 (Table 1). For the former, redundant sequences were not filtered out. For the latter two, sequences were clustered at 40 and 20% sequence identity, respectively, and only cluster representatives are kept. Thus, CATH should be easiest and CATH S20 most difficult among the three datasets.

**Table 1.**
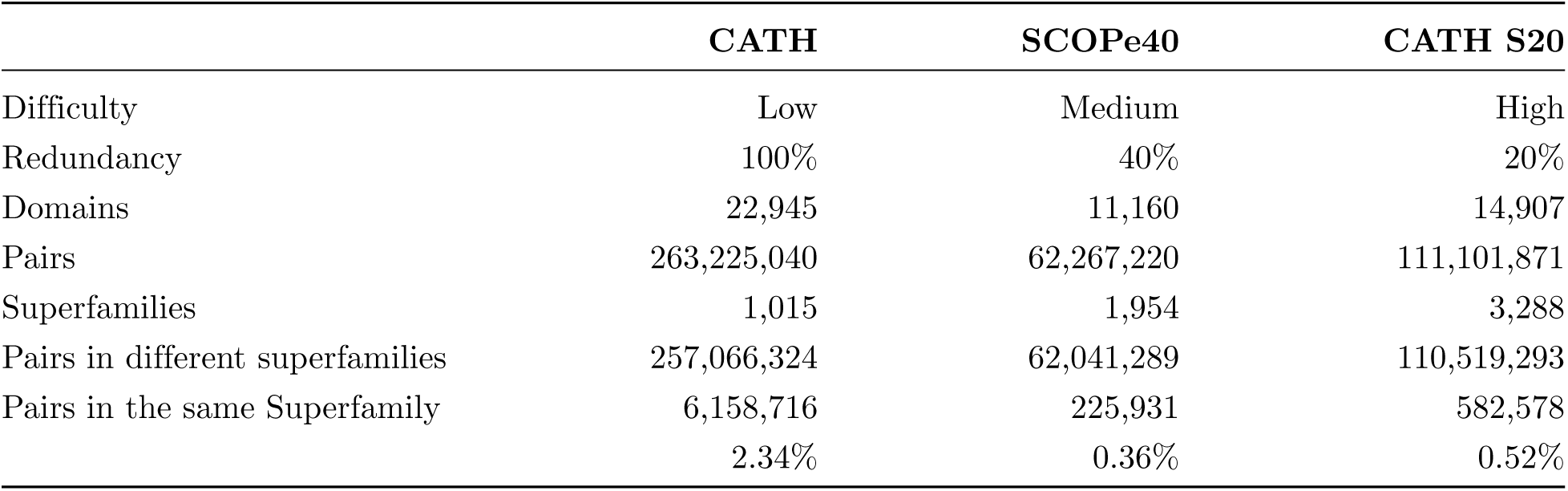
Overview of CATH and SCOPe40 datasets, including redundancy levels and pair counts.

To achieve a fair comparison, it is important that datasets are large with little selection bias and that the ratio between positive and negative pairs is small, as it would be the case for a realistic RHD task. The datasets cover over 1,000 superfamilies each and more than 10,000 individual domains. The number of domain pairs ranges from 62 to 263 million, with the number of pairs within the same superfamily being a small fraction of all pairs (0.36% via 0.52% to 2.34%, see Table 1). The greater difficulty of the non-redundant sets is also reflected in 4.5 - 6.5-fold lower percentage of positive pairs.

Given the three datasets, we compared the performance of six different methods as summarised in Table 2. We measured performance using AUC and AUROC1 to make results comparable and independent of individual scoring schemes of the method. While AUC provides a global measure of separability between positive and negative pairs, it does not necessarily reflect practical performance in our setting. Because it integrates over all possible score thresholds, AUC can remain high even when a method fails to rank true homologues among the top hits, particularly in datasets with strong class imbalance where true negatives dominate. In contrast, the AUROC1 metric captures a more application-oriented perspective by quantifying the sensitivity up to the first false positive for each query. Consider a biologist’s common use case of searching a sequence against a large database of hundreds of millions of proteins. In a database search returning millions of candidates, a biologist will only ever examine the top-ranked hits. Note that in line with [11] AUROC1 considers pairs with a different fold a false positive, while AUC considers a pair with different superfamily already a negative pair.

**Table 2.** Performance (AUC and AUROC1) comparison of different methods on the redundant (CATH), 40% redundant (SCOPe40), and 20% redundant (CATH S20) datasets. The AUC score captures the method’s global separability between positive and negative pairs, while AUROC1 quantifies the sensitivity up to the first false positive for each query.

| Repr. | Method | Ext.? | CATH |  | SCOPe40 |  | CATH S20 |  |
| --- | --- | --- | --- | --- | --- | --- | --- | --- |
|  |  |  | AUC | AUROC1 | AUC | AUROC1 | AUC | AUROC1 |
| Seq, HHM profiles | HHblits | yes | 0.92 | 0.70 | 0.94 | 0.64 | 0.75 | 0.41 |
| Seq, 20-letter 3Di | Levenshtein | no | 0.90 | 0.56 | 0.89 | 0.33 | 0.82 | 0.16 |
| Seq, Q8 SS | Levenshtein | no | 0.95 | 0.50 | 0.90 | 0.28 | 0.81 | 0.05 |
| Seq, Q3 SS | Levenshtein | no | 0.95 | 0.49 | 0.89 | 0.24 | 0.81 | 0.08 |
| Seq, Q3 SS pred. | Levenshtein | no | – | – | – | – | 0.81 | 0.28 |
| Str, Foldseek | FS-pair | no | 0.97 | 0.72 | 0.95 | 0.62 | 0.77 | 0.39 |
| Str, 3D coordinates | US-align | no | 0.99 | 0.80 | 0.96 | 0.67 | 0.90 | 0.49 |

Standard AUC rewards ranking true positives even e.g. at rank 1000 or later, since it is still ahead of hundreds of millions of non-relevant proteins. AUROC1 only considers relevant hits up to the first non-relevant hit. This way it reflects how effectively a method retrieves true homologues before introducing erroneous hits — an aspect that is crucial for realistic database search scenarios. However, AUROC1, by design, ignores performance beyond the first false positive and therefore underrepresents methods that maintain good ranking performance deeper into the hit list. Therefore, we also considered the individual scoring distributions, split for positive and negative pairs, to gain insight into how well methods separate positive pairs from the negative pairs (see S3 Note for AUROC1 scores and scoring distributions). Together, these three metrics provide complementary insights: AUC summarises overall discriminative power, while AUROC1 highlights early precision and retrieval quality in the high-confidence range, and the score distributions show how well each method separates positives from negatives across the full range.

We now want to evaluate the structural alphabets on this fold recognition benchmark and put them into comparison with basic sequence alignment tools as well as gold standards in the field, which are structural alignment and HMMs.

### 3.2 Full 3D structural alignment, Foldseek and HMMs are gold standards

3D coordinate-based alignment methods directly compare atomic-level structures. Structural alignments were computed with US-Align [8], which reports a template modelling score (TM-score). An advantage of TM-scores is that they are normalised between 0 (poor) and 1 (good), where values above 0.5 indicate shared fold-level structural similarity. [22]. On the CATH dataset, structural alignment achieves a nearly perfect AUC score of 99%.

Interestingly, structure alignment does not perform perfectly, in particular on the more difficult datasets. One reason may be structural flexibility, another reason inconsistencies in the superfamily classification. For example, we observed instances where highly similar structural folds were classified into different SCOPe superfamilies (see Fig. 2). Visual inspection confirmed structural similarity, suggesting that these classifications into different superfamilies may stem from functional differences or a lack of evidence for homology during manual curation (see Uncertain database labels). Such cases highlight the inherent ambiguity in superfamily definitions and the challenges of achieving perfect classification.

**Fig 2.**
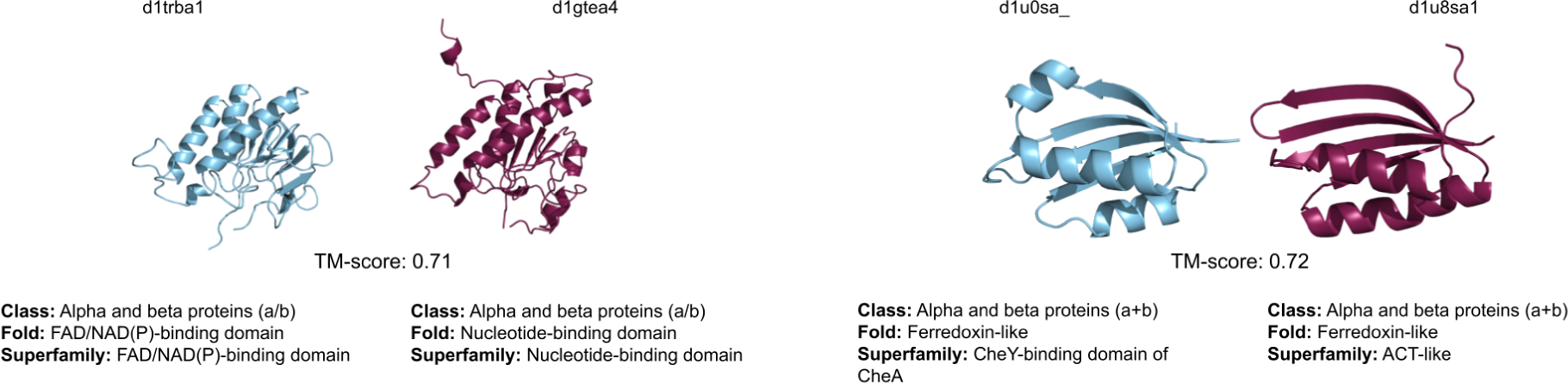
Examples of false positives of TM-score classification. **Left:** Classification in a different fold and different superfamily. Visually, both structures look very similar, which is also evident from the high TM-score. They are, however, not only in different SCOPe superfamilies but even different folds, sharing only the class-level classification. **Right:** Classification in the same fold but different superfamily. Upon visual inspection, the beta sheet motif and the two alpha helices clearly show a strong resemblance.

As a first result and in line with previous studies, the Hidden Markov Model approach HHblits successfully retains a strong signal from primary structure in remote homology detection (RHD). HMMs capture a statistical signature of a sequence family, making them more effective for RHD than basic sequence comparison. However, notably the captured signal is less pronounced on the most challenging CATH S20 dataset and more substantial for SCOPe40 and CATH, which provide more of the data needed to train the HMM.

It is important to note that by including HHblits we veer off from a pure representation comparison, and compare representations to a practical tool. This was done to not underestimate the information amino acid sequences hold and reflect the state of the art usage of those. However HHblits should be considered somewhat outside the formal competition, as it leverages external data: unlike other methods evaluated here, HHblits condenses the information contained in UniRef30, a database comprising over 30 million protein sequences. This gives HHblits a substantial edge, as it effectively exploits the ever-growing wealth of sequence data. Overall, HHblits achieves an AUC of 92% on the CATH dataset.

In fact, this principle of learning from massive sequence databases culminates in the recent introduction of protein embeddings—statistical sequence models trained on billions of sequences [20].

### 3.3 Non-redundant datasets are consistently more challenging

The more non-redundant a dataset is, the fewer “easy” remote homologues exist, making the dataset more challenging. As Table 2 shows, this consistently applies to all methods, including structural alignments. It is expected for sequence-based methods but surprising for structural methods. For these there may be three main reasons, namely variability of protein structures, imbalance of the classification tasks, and manually curated superfamily labeling beyond structural similarity in the domain annotations (see Uncertain database labels).

Interestingly, there is also a pronounced difference in difficulty for the 20% and 40% redundancy datasets. This may be explained by a transition at around 30% sequence identity, the so-called twilight zone for sequence comparison.

There is, however, also one outlier to this trend, where HHblits performs better on SCOPe40 than on the redundant CATH dataset and also better than the structural alphabets.

### 3.4 Fold information is retained across structural alphabets

The main result of this article is that across all benchmarks, structural alphabets perform closely behind the US-align reference, with the 20-letter 3Di alphabet being 7-9% behind US-align and retention of the fold-recognition signal being 32-40% above chance. The Q8 and Q3 alphabet are 4-9% behind US-align while retaining the fold-recognition signal is 31-45% above chance. For the difficult CATH S20 dataset, which resembles most closely the remote homology problem setting, the 3Di alphabet comes close to the gold standard structural alignments in terms of AUC (see Table 2 and Fig. 3), achieving 91% of US-align’s performance and showing scoring distributions akin to US-align (see Fig. 4, S3 Note).

**Fig 3.**
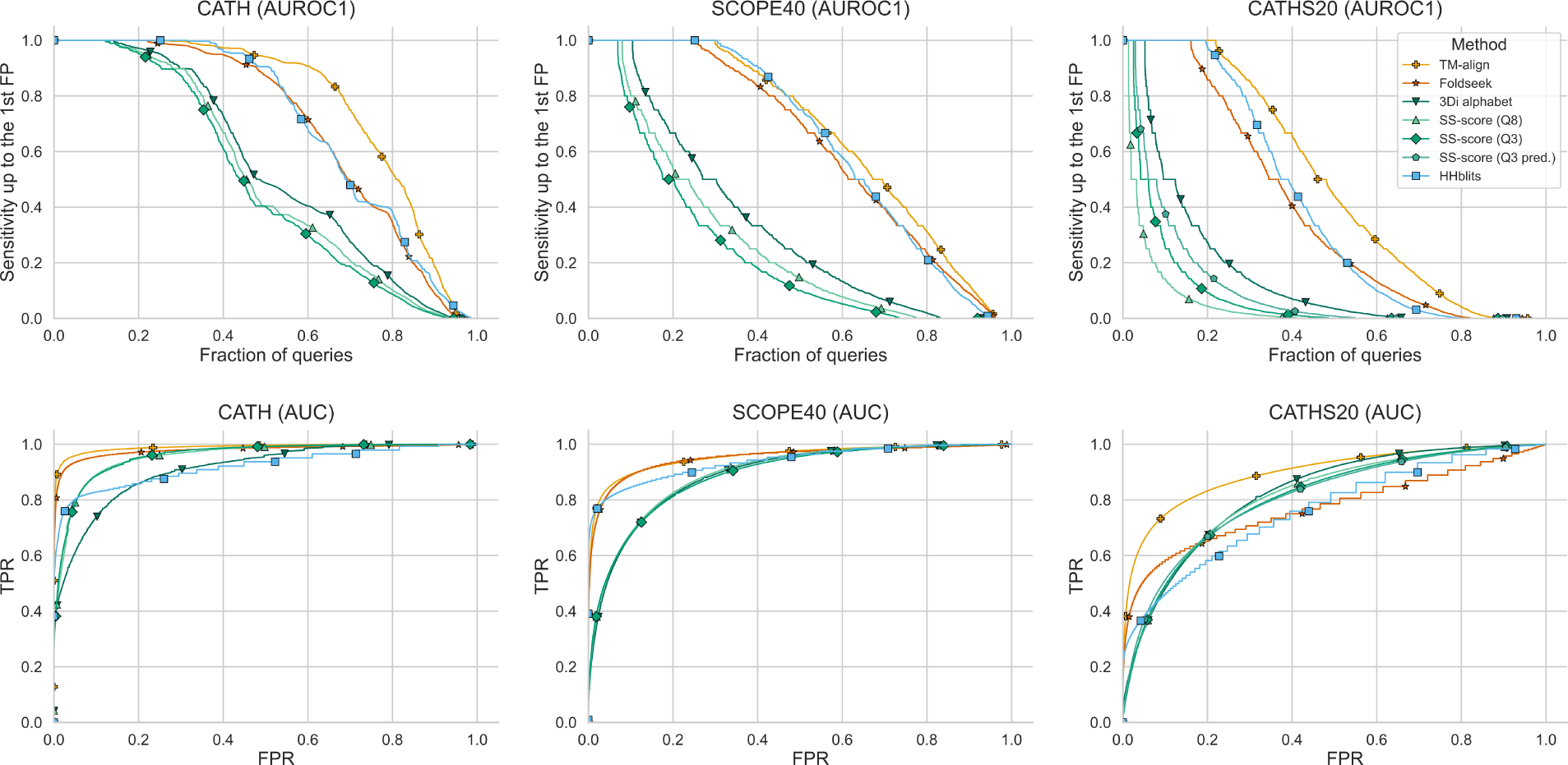
AUROC1 and AUC curves for the three datasets of varying difficulty. **Top:** AUROC1 plots show cumulative distribution of sensitivity up to the first false positive (FP). True positives (TPs) are matches of the same superfamily, false positives (FPs) of different fold. Methods were grouped by color, amino acid alphabet (blue), structure alphabet (green), and structure alignment (yellow). Here again US-align outperforms all other methods, while the structure alphabet-based methods perform similarly well. We observe that datasets with less redundancy are across all methods, consistently more difficult to classify. **Bottom:** AUC curves of false positive rate (FPR) against true positive rate (TPR). The corresponding values are in Table 2. Structure-based alphabets perform similarly well and generally outperform the sequence-based HHblits. Structure alignment consistently outperforms all other methods and can be considered the gold standard. For the medium-difficulty dataset, SCOPe40, HHblits performs better than the structure-based alphabets; however, this could be due to SCOPe superfamily assignment employing an HMM search.

**Fig 4.**
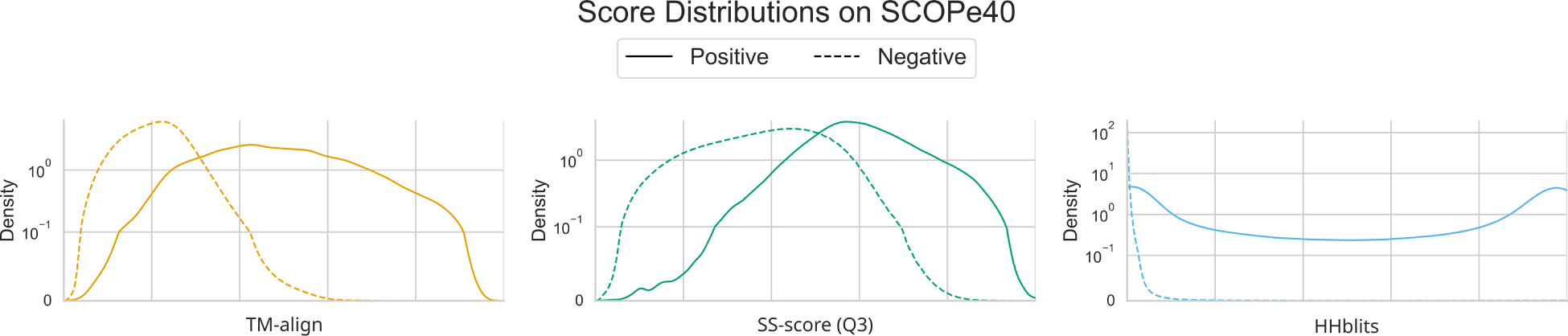
Score distributions of TM-align, Q3 alphabet and HHblits on SCOPe40 dataset; continuous line shows distributions for pairs in the same superfamily, dashed line for those that are in different superfamilies. This allows us to see how well methods separate positive pairs from negative pairs. While US-align and the Q3 alphabet are rather unimodal distributions, HHblits shows a bimodal distribution for the positive pairs, with roughly half of the pairs being scored at 100 % probability of being homologous and the other half at around 0% probability, meaning that in a practical scenario HHblits categorises half of the positives as false negatives, even at the easiest difficulty (CATH, see S3 Note). Note that distributions have been normalised per group, i.e., for positive pairs and negative pairs each per method and dataset; also, the y-axis is symlog scale. This is due to the imbalance in the dataset. For score distributions of all methods on all datasets see S3 Note.

### 3.5 Three-state secondary structure retains most of the fold-recognition signal

Surprisingly, the same holds true for the much simpler Q3 alphabet. In fact, for the redundant CATH dataset, the Q3 alphabet even outperforms 3Di and across all benchmarks retains 96%, 93% and 90% of the US-align signal (easiest to most difficult, see Table 2 and Fig. 3). This is remarkable, since in contrast to 3Di, secondary structure abstracts 3D structure by clustering and labelling combinations of phi and psi angles derived from atomic coordinates. This means despite missing explicit spatial atomic distances, orientations and tertiary contacts, the Q3 representation is still capable of retaining most of the fold-recognition signal.

### 3.6 Increasing alphabet granularity provides limited additional fold information

In addition to the Q3 alphabet, we also benchmarked the Q8 structural alphabet, offering another level of granularity: while Q3 represents a coarse 3-state alphabet (helix, strand, loop), Q8 offers a granular 8-state alphabet (differentiating helix types, strands, and specialised turns). As Table 2 shows, the results for both representations are nearly identical. I.e., an increased granularity does not translate to increased performance. This is because the increased granularity concerns only a minority of residues (see S4 Note) and unlike amino acid alphabets, where large differences in frequencies can be attributed to physicochemical properties (e.g., the rarity of cysteines forming disulphide bonds), such specific roles are not ascribed to the more detailed secondary structure descriptions of the Q8 alphabet. To further investigate this, we calculated the Shannon entropy *H* of Q3 and Q8 alphabets on the SCOPe40 dataset (see Table 3). Here we found that while the Q3 alphabet reaches close to maximum possible entropy, Q8 does not. The reduced normalized entropy at Q8 level (80.7% vs. 96.2% at Q3) supports our notion that several Q8 states occur with very low frequency in the dataset. Q8 represents 8 states, but statistically behaves closer to a 5-state alphabet. Since standard accuracy metrics are frequency-weighted, rare states contribute negligibly to reported performance.

**Table 3.** Comparison of Shannon entropy and effective state utilization between Q3 and Q8 alphabet on SCOPe40. Effective states are computed as 2*^H^*, where *H* is the observed Shannon entropy. The reduced alphabet utilization at Q8 level (66.8% vs. 95.7%) demonstrates that the perceived gain in granularity is not reflected in the data.

| Alphabet | Maximum possible entropy | observed entropy H | normalized entropy | Effective states | Alphabet utilization |
| --- | --- | --- | --- | --- | --- |
| Q3 | 1.58 bits | 1.52 bits | 96.2% | 2.87 | 95.7% |
| Q8 | 3 bits | 2.42 bits | 80.7% | 5.34 | 66.8% |

Regarding an improved secondary structure representation, we wanted to determine whether we can expand the secondary structure information encoded in Q3 by injecting residue contacts to align them further with 3Di. In a preliminary experiment we extended the 3-letter alphabet to 6 letters representing helix, strand, and turn with and without a long-range contact [37]. Against our expectations, this did not substantially improve performance. Overall, we achieved an AUC of 0.90 for the 40% redundant SCOPe40 dataset, which is virtually on the same level as regular secondary structure (0.89).

### 3.7 Q3 and 3Di encode overlapping structural information

As a reminder, SS and 3Di are conceptually distinct: SS captures the phi-psi angles of residues, whereas 3Di describes the local structural environment surrounding each residue. 3Di uses the 20 letters

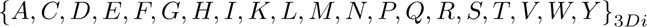

To understand why secondary structure performs as well as 3Di, we looked at the correlation of representations against each other. For this, we picked the CATH dataset, as it is the one containing the most entries. We find that certain 3Di characters strongly correlate with very specific secondary structure elements (SSEs). The characters *{V, L, S, C}*_3_*_Di_* all show a Pearson correlation coefficient greater than 0.75 for helices and a negative correlation less than 0.4 for strands (see Fig. 5A). Moreover, it is possible to identify four groups within the 3Di alphabet that correlate between moderate and very strong with SSEs: *{V, L, S, C, N }*_3_*_Di_* with helix (and negatively with strand), *{I, W, K, T, F, M, G}*_3_*_Di_* with strand (and negatively with helix), *{A, H, R}*_3_*_Di_* with loop, and the rest, *{D, Q, P, Y, E}*_3_*_Di_*, being wildcards of either strand or loop or all three. Particularly interesting is that over 40% of the dataset is comprised of just 3 of these letters (*V, D, P*) and over 60% by just 7 letters (see Fig. 5B). The letters *V, L, S* make up 17.7, 5.5 and 5.2% respectively, while being very strongly (*>* 0.9) correlated with helices. Our findings therefore overall suggest that the Q3 and 3Di structural representation overlap substantially but are not equivalent.

**Fig 5.**
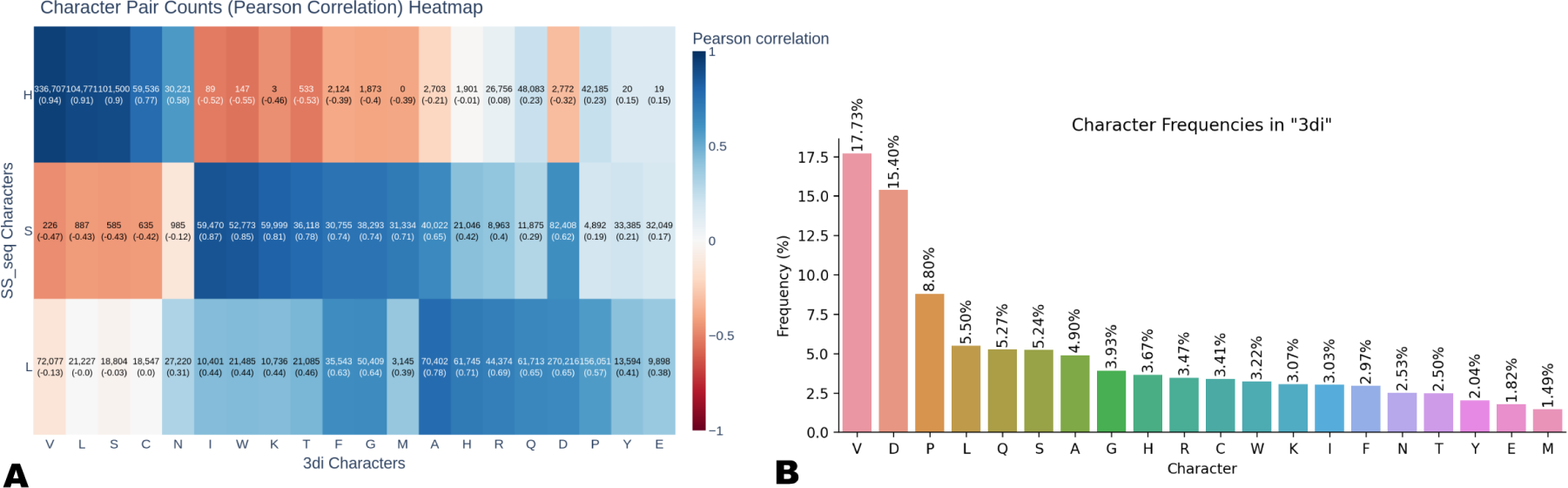
**A**: Heatmap of correlation between Q3 secondary structure and 3Di representation. Cells are annotated by total occurrence of this pairing, Pearson correlation in parentheses. Colour according to Pearson correlation. Most of the 3Di letters correlate strongly with a specific SS element, particularly the correlation for *V, L, S,* with helix or *I, W* with strand, suggesting SS information is implicitly encoded in the 3Di representation. **B**: Distribution of 3Di characters in the CATH dataset. Over 40% of the dataset comprises just 3 out of the 20 letters, and over 60% just 7 out of the 20 letters of the 3Di alphabet.

Additionally, some 3Di characters show moderate to strong correlation with particular amino acids, e.g., *{V, L, S}*_3_*_Di_* with alanine and leucine or *A*_3_*_Di_* with glycine, serine, and threonine (see S5 Note). Conversely, glycine is moderately to strongly correlated with *{A, H, Q, R}*_3_*_Di_*, suggesting that 3Di carries some primary information as well. These observations are broadly consistent with prior work. The strong helix association of *V*_3_*_Di_* corroborates both Heinzinger et al. [38] and Pantolini et al. [39], who similarly identify *V*_3_*_Di_* as the dominant helix token. For strand-associated states, our grouping agrees with Heinzinger et al. for *W*_3_*_Di_*, but diverges for *A*_3_*_Di_* and *Y*_3_*_Di_*, which in our correlation analysis associate with loop and show wildcard behaviour, respectively, rather than strands. Similarly, while both Heinzinger et al. and Pantolini et al. assign *D*_3_*_Di_* predominantly to loops, our analysis places *D*_3_*_Di_* among the wildcard states with no strong exclusive SSE association. This discrepancy may reflect differences in dataset composition or the quantitative versus visual nature of the two assessments.

### 3.8 Sequence-predicted Q3 preserves the global fold-recognition signal

In order to test the viability in a setting with no structure available, we benchmarked predicted Q3 strings on the difficult CATH S20 dataset. Here we found that the performance for predicted Q3 strings holds up to those determined from structure and even shows substantially improved performance in terms of AUROC1 (see Table 2, Fig. 3).

### 3.9 Global signal retention does not imply equivalent early retrieval

In terms of AUROC1, US-Align, HHblits and Foldseek clearly outperform all other methods (see Fig. 3): US-Align due to its exhaustive structural information, HHblits due to its access to the vast UniRef30 sequence database, and Foldseek by combining structural alphabet encoding with an optimised search pipeline and rigorous significance cutoffs.

US-Align stands out as one of the top-performing methods with AUROC1 values of 0.80, 0.67 and 0.49 for the three difficulty benchmarks, which is expected given its exhaustive access to structural information.

A similar picture emerges for Foldseek, which achieves competitive AUROC1 scores across all three datasets (0.72, 0.62, 0.39), approaching the gold standards despite relying on the 3Di structural alphabet rather than full coordinate-based alignment.

While it may appear that HHblits comes even closer to US-Align here than Foldseek, achieving AUROC1 scores of 0.70, 0.64 and 0.41, it is important to note that AUROC1 provides only a limited view and not the full picture (as evident from the AUC plots and AUC table). For a more complete insight, we examined the score distributions (see Fig. 4, S3 Note). This analysis reveals that while US-Align and Foldseek generally assign higher scores to positive pairs than to negative pairs, HHblits shows a bimodal behaviour: for roughly half of the positive pairs, it is absolutely certain they are homologous; however, for the other half, it confidently misclassifies them as non-homologous, assigning scores close to 0. This results in approximately half of the positives being classified as false negatives.

The fact that US-align, Foldseek and HHblits remain much stronger for early retrieval should not come as a surprise, however. These are contextual systems and should not be considered in the same ballpark as the controlled alphabet comparisons we presented.

### 3.10 Q3 prioritises structural matches in an illustrative newt case study

Encouraged by the overall results, we wanted to demonstrate the use of the Q3 secondary structure alphabet in function annotation based on remote homologues with an illustrative use-case.

The highest quality of annotations can be expected from direct, experimental evidence. By its very nature, such evidence is biased towards established model organisms. [40] analysed annotations from 12 model and 241,860 non-model organisms and found that out of a total of 75,304,476 annotations, less than 1% are direct, experimental evidence and only 0.03% are direct, experimental evidence for non-model organisms. Therefore, accurate computational inference of annotations across vast evolutionary distances is essential.

As an illustrative example, we consider the Iberian ribbed newt, whose genome was very recently sequenced [29] and hence there is no direct evidence for any of its gene products. Its closest relative, axolotl, is intensely studied but not yet well annotated, either. The closest amphibian, Xenopus, and well-annotated chordates like zebrafish and mouse are evolutionary distant. To put this into numbers: There are over 250 million unreviewed sequences in UniProt [31] and less than 600,000 reviewed ones. Bacteria is the largest kingdom for both. Within eukaryotes, fungi, including yeast, is largest. Regarding newt, axolotl is not among the top 250 species in terms of number of reviewed protein sequences, and xenopus and zebrafish rank only in positions 14 and 15 (see uniprot.org/uniprotkb/statistics). To summarise, newt protein sequences are highly challenging to annotate.

From the Iberian ribbed newt genome sequence we derived two sets of protein sequences of size 99 and 20, respectively. The former is a random selection from 33,854 newt candidate genes, which match a protein-coding region in another organism at a significant E-value [29]. For the latter, such a match does not exist, which implies that they are the most challenging to annotate. We limited the sizes of both sets, as structure prediction and pairwise structural alignments were computationally very demanding and supercomputing resources were limited.

The 99 and 20 newt proteins were compared against 496,866 proteins from Swiss-Prot. Swiss-Prot was chosen instead of the entire UniProt database, as Swiss-Prot entries are manually reviewed and contain annotations. Swiss-Prot entries were limited to those for which a predicted AlphaFold structure of high quality existed.

With benchmark and library defined, we have to deal with comparing domains with proteins. The Levenshtein algorithm used so far compares entire sequences. This is adequate for domains, but not for proteins, for which a local alignment is needed. Hence, we introduced a further improvement for secondary structure: the alignment algorithm. In particular, we wanted to determine whether the results can be improved by moving from a global to a local alignment and from a simple match score to a secondary structure substitution matrix. We implemented local alignments with the Smith-Waterman algorithm [34]. This entailed the definition of gap penalties and a substitution matrix. For the gap penalties, we experimented with values mirroring the default BLAST settings [21] and found a penalty of -11 for opening a gap and -3 for extending a gap as optimal. The construction of the substitution matrix followed a similar procedure as BLOSUM [27], the details are described in Materials and methods.

As the newt proteins are not annotated, there is no ground truth. Instead we relied on the TM-score from structural alignment as indication for homology. More precisely, following Xu and Zhang [35] TM-score above 0.5 corresponds to substantial structural similarity and similar to Liu et al. [41], lacking a proper ground truth we considered protein pairs with TM-score above 0.5 to be “homologues”. As a lower bound reference, we chose BLAST and PSI-BLAST, as they were used by [29] in their initial annotation process.

Table 4 summarises the results for the nearly 50 million comparisons. BLAST and PSI-BLAST achieve an AUC of 0.51 and 0.52. Structural alignment has, by definition, an AUC of 1.0, since it is not an independent method performance but the reference used to construct the proxy labels. The key result is that secondary structure performs close to structural alignment with an AUC of 0.94.

**Table 4.** Performance (AUCs) comparison of different methods on proteins from Newt.

| Structure | Methods | AUC |
| --- | --- | --- |
| Seq, amino acid | BLAST | 0.51 |
| Seq, amino acid | PSI-BLAST | 0.52 |
| Seq, Q3 SS | Smith-Waterman | 0.94 |
| Str, 3D coordinates | US-Align | 1.0 |

For the second function annotation task we considered more challenging newt proteins, for which Brown et al. could not find any related sequences. Here we want to emphasise, that we want to present this as an illustrative example. This is not a validated functional annotation of unannotated newt proteins. We applied the Smith-Waterman algorithm and observed that overall the scores for pairs that were labeled homologous achieved higher scores than non-homologous pairs (see Fig. 6b). We then evaluated the 5 proteins with the best scores. For 7 out of these 20, we could find hits, which can be considered homologues by TM-score.

**Fig 6.**
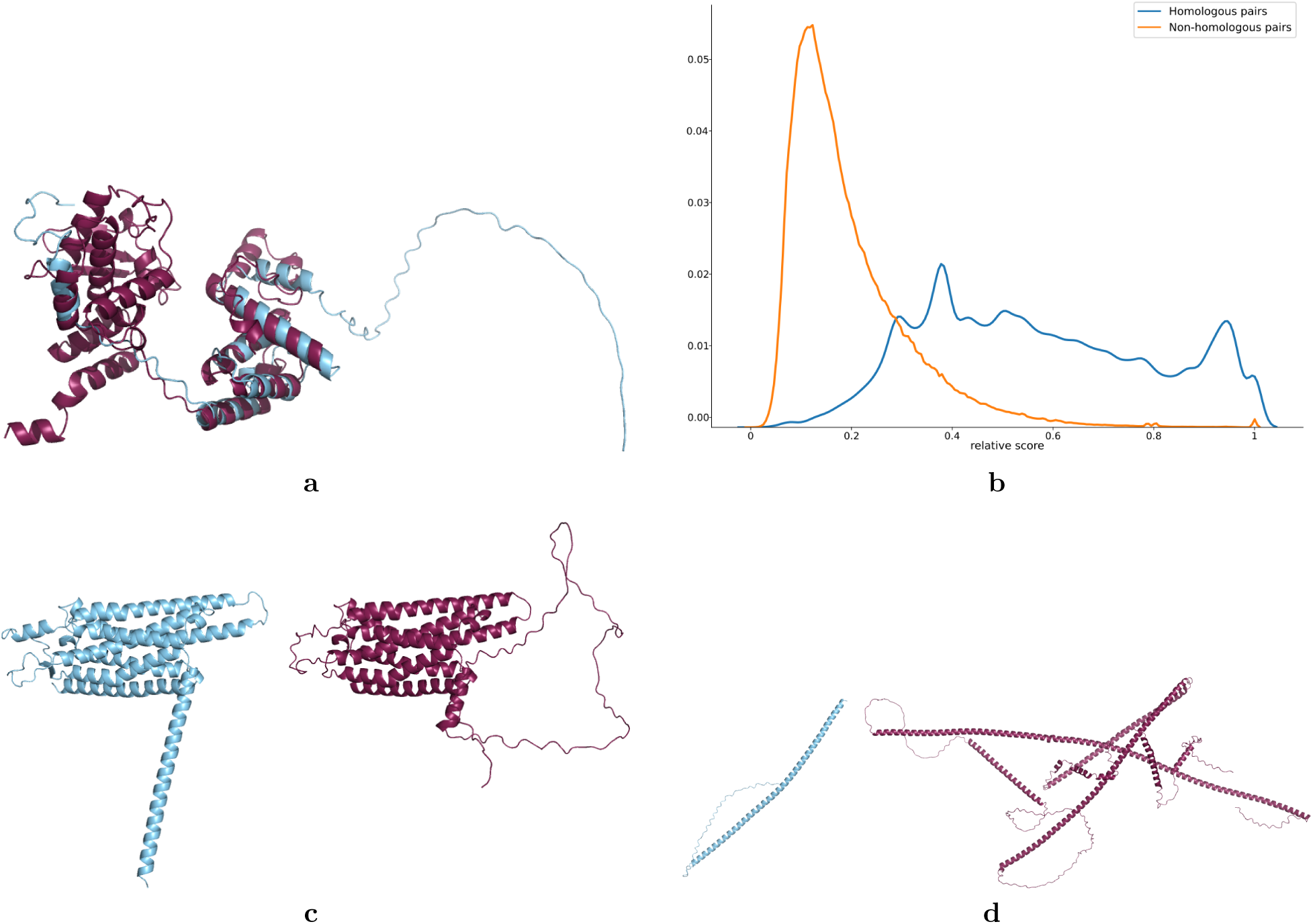
a: A query for a newt protein mRNA116414 (blue) produces the protein O50655 (magenta) using secondary structure comparison only. Structural alignment supports the strong resemblance and possible remote homology. **b:** The distribution of the relative scores for homologous and non-homologous pairs. The relative scores are as described in Materials and methods. The distributions for the homologous and non-homologous pairs were calculated individually, so the reported density functions only apply to part of all obtained scores. The distributions for homologous pairs are coloured blue; the distributions for non-homologous pairs are coloured orange. **c–d:** Two more examples of query (blue) side-by-side with the top hit (magenta). The left demonstrates a good example with strong structural similarity, while right shows a caveat of the SS strings, as they are prone to match long helical motifs that are unrelated.

Given the structural similarity between *mRNA116414* and *O50655*, the unannotated protein from newt is likely an integrase/recombinase (see Fig. 6a). As a comparison, we also ran PSI-BLAST for these 20 newt proteins against our library. In 15 out of 20 cases, the Smith-Waterman TM-scores were better than the PSI-BLAST ones (see S2 Table). Two more examples with their best match are shown in Fig. 6c, 6d, showcasing both the potential and caveats of the approach. For a side-by-side of all 20 queries with their best match from the SW method, see S2 Fig.

Xu and Zhang [35] argued that a TM-score of 0.5 is suitable to indicate the same fold, hinting at possible homology. To define a similar threshold for secondary structure, we pursued a Bayesian approach and calculated the posterior probabilities of protein pairs belonging to the same superfamily based on their secondary structure alignment scores, providing a probabilistic framework for interpreting these scores. We found that a score of 0.85 is suitable as a heuristic threshold, which could be used in practice to deploy our algorithm in future work (see S6 Note), for a fully operational cutoff, statistical analyses on a larger set will be necessary.

## 4 Discussion

### 4.1 What extreme structural compression reveals about protein folds

As a central finding of our analysis, we demonstrated that much of the fold-recognition signal remains even after explicit distances, orientations and tertiary contacts are removed. The Q3 alphabet still retains at its best up to 96% of the signal that US-align captures. This is a striking result: most of the global fold-recognition signal (measured by AUC) survives without atomic coordinates, torsion angles and spatial contacts. This geometric detail appears to account for the remaining margin of 4-9% in AUC. Meanwhile, what remains after the compression to Q3, is the order and organization of secondary structure elements, the sequence in which helices, strands and loops occur along the chain. Our results suggest that this ordering, rather than the precise 3D geometry, carries a large share of the discriminative signal used in fold recognition.

This interpretation is consistent with how secondary structure has historically been treated as an intermediate abstraction between primary sequence and tertiary fold. Assignment methods such as DSSP [24] and STRIDE [42, 43] already reduce atomic-level geometry (backbone hydrogen bonds, torsional angles) to a small discrete alphabet — eight states (Q8) or, more coarsely, three (Q3: helix, strand, coil). That such a coarse-grained, order-preserving abstraction suffices to capture 90%-96% of the 3D structure alignment signal indicates that a substantial part of the fold-level signal carried by the topological order and arrangement of secondary structure elements rather than by the fine geometric detail these methods discard.

Recent embedding-based approaches reinforce this view from a different angle. Methods that learn secondary-structure or fold representations directly from sequence — Dallago et al. [44], Elnaggar et al. [20], extending earlier PSIPRED [45], S4PRED [46], and SSE-PSSM [47] approaches — achieve accurate structural characterization without ever constructing an explicit 3D model. TM-vec/DeepBLAST [48] goes further, predicting TM-scores directly from sequence embeddings and replacing costly pairwise structural alignment with embedding similarity. That these compressed, geometry-free representations perform well at fold-level tasks is further evidence that our findings are not an isolated artifact of our specific setup, but reflect a broader property of how fold information is distributed across levels of structural abstraction.

### 4.2 Alphabet size is an incomplete measure of structural information

A further finding of our analysis is that the number of nominal states an alphabet uses does not predict how much fold-discriminative information it carries. To compare alphabets on equal footing, we paired Q3, Q8, and 3Di with the same simple alignment algorithm, Levenshtein distance, rather than an alphabet’s native, more elaborate scoring scheme: isolating what each alphabet encodes from how a full search system exploits it. Under this comparison, Q8 achieves essentially the same AUC as Q3 across all three benchmarks (CATH: 0.95 vs. 0.95; SCOPe40: 0.90 vs. 0.89; CATH S20: 0.81 vs. 0.81), despite offering more than double the number of nominal states. The 20-state 3Di alphabet fares only marginally better on CATH S20 (0.82) and is on par with or below Q3 and Q8 on the other two datasets (Table 2).

An entropy analysis of the secondary-structure alphabets clarifies why extra states fail to translate into extra information. Q3 utilizes 95.7% of its theoretical alphabet capacity (normalized entropy: 96.2%; effective states: 2.87 of 3 possible), meaning its three states are used close to maximally. Q8, by contrast, utilizes only 66.8% of its capacity (normalized entropy: 80.7%; effective states: 5.34 of 8 possible; see Table 3), meaning that Q8 behaves statistically like a 5 or 6-state alphabet, because several of its states occur rarely, rather than each contributing an independent axis of information.

This redundancy is not unique to Q8. Mapping the 20-letter 3Di alphabet against the coarser Q3 categories shows strong correlations between individual 3Di characters and a single Q3 state, several 3Di letters correlate with the helix category at Pearson coefficients above 0.9 (see Fig. 5A), indicating that much of 3Di’s apparent granularity subdivides categories Q3 already captures, rather than encoding structurally independent information. The 3Di alphabet is also used highly unevenly: its three most frequent characters (V, D, P) together account for 41.9% of all occurrences, while its least frequent characters occur in under 2% of positions (see Fig. 5B). Rare, redundant, and correlated states of this kind explain why increasing the alphabet size does not produce a proportional gain in fold-recognition performance.

This is consistent with recent work on structural alphabet design: while larger alphabets have been argued to improve retrieval performance [49], our results suggest such gains diminish once a coarser encoding already captures the dominant structural signal. Zerefa et al. [50] offer a complementary data point: combining their 3Dn alphabet (backbone dihedral angles) with 3Di improves retrieval over either alone, because the two alphabets capture geometrically distinct properties. Read together with our result, this points to the same principle from opposite directions: additional states help only when they capture a genuinely orthogonal structural property, not merely when they subdivide an alphabet more finely.

### 4.3 Representation fidelity and search performance answer different questions

The comparisons in the previous section measured what an alphabet encodes, using a single, minimal alignment algorithm applied uniformly across alphabets. This isolates representation fidelity, but it discards nearly everything an actual use-case search tool adds on top: substitution matrices tuned to the alphabet’s statistics, local rather than global alignment, statistical calibration (E-values), profile information drawn from external databases, and indexing for large-scale search. These two questions, how much fold information a representation carries, and how well a complete system retrieves true hits near the top of a ranked list, turn out to have different answers.

By AUC, the plain alphabets are close to the engineered systems. By AUROC1, which reflects whether true positives are ranked consistently at the very top of the hit list, the metric that matters for practical database search, the gap is large: Levenshtein-based Q3,Q8 and 3Di top out at 0.49–0.56 on CATH and fall to 0.05–0.16 (with exception for the predicted Q3) on the more redundancy-controlled CATH S20, while Foldseek, HHblits, and US-align all remain in the 0.39–0.80 range across all three benchmarks (see Table 2). Representation fidelity and early-retrieval performance are therefore not the same axis: a coarse alphabet can capture most of the discriminative signal an alignment task requires while still ranking true hits far less reliably than a calibrated search system does.

HHblits makes this point sharply. Despite using an entirely non-structural representation, namely sequence profiles built iteratively from an external database, HHblits achieves the second-highest AUROC1 of all methods we tested for SCOPe40 and CATH S20 and the third-highest for CATH, exceeding even Foldseek’s structure-based pipeline on two benchmarks (SCOPe40: 0.64 vs. 0.62; CATH S20: 0.41 vs. 0.39). This is a striking outcome given that HHblits’ overall AUC trails structural methods, particularly on CATH S20 (0.75 vs. 0.90 for US-align). The explanation is visible in the score distributions (see S3 Note): HHblits assigns roughly half of true positives scores near zero, a substantial loss in raw discriminative content, yet the positives it does score highly are ranked with enough confidence, via its calibrated profile search, to dominate the top of the list. Our Rad52 example (see Fig. 1) illustrates the failure mode directly: HHblits finds almost no similarity between two structurally near-identical proteins via profile alignment, while secondary structure alignment recovers the shared motif immediately — yet in aggregate, HHblits’ calibration and external-database machinery still deliver strong early retrieval elsewhere.

Foldseek shows the complementary case: its full pipeline, the 3Di alphabet together with amino acid sequence plus a customized substitution matrix, local alignment, and E-values, roughly doubles AUROC1 over the same 3Di alphabet under plain Levenshtein alignment (e.g., SCOPe40: 0.62 vs. 0.33), despite AUC moving only modestly (0.89 and 0.95 on SCOPe40). The combined representation of 3Di and AA is and especially the algorithmic engineering built around it is what drives the early-retrieval gain.

### 4.4 Sequence prediction broadens the possible use of Q3

Our benchmark so far relied on secondary structure derived from experimentally determined or predicted structures, however the most consequential use case is one where no structure exists at all. We therefore tested predicted Q3 strings, generated directly from sequence, on the CATH S20 benchmark. Predicted Q3 performs essentially on par with structure-derived Q3 (AUC: 0.81 for both) and, notably, shows higher AUROC1 (0.28 vs. 0.08), showing that early retrieval is not degraded by prediction error and may even benefit from it, though a single dataset is too little evidence to treat that AUROC1 gain as a real effect rather than noise.

This robustness is consistent with the known error profile of secondary structure prediction. Assignment itself has an inherent accuracy ceiling of approximately 88–92%, driven by “chameleon” regions of up to 11 residues that adopt different states depending on context, and experimentally determined structures already disagree by 5–15% between X-ray and NMR models [51]. State-of-the-art prediction methods now reach 81–89% accuracy [20, 38, 52], close enough to this natural ceiling that the residual prediction error apparently does not meaningfully erode the fold-recognition signal Q3 carries.

This result is encouraging for the many practical settings, such as newly sequenced genomes, proteins without solved or confidently predicted structures, where Q3 must be derived from sequence alone. However, we tested predicted Q3 on a single benchmark (CATH S20); broader benchmarking across CATH and SCOPe40, together with runtime measurements comparing prediction-based and structure-based pipelines, is still required before this can be treated as a general result.

### 4.5 Structural similarity enables hypothesis generation, not functional assignment

To illustrate Q3’s practical use when no annotation exists, we applied it to the Iberian ribbed newt, a recently sequenced genome with no direct experimental evidence for any gene product [29]. This should simulate a situation typical of the vast majority of sequenced life: fewer than 1% of all UniProt annotations rest on direct experimental evidence, and only 0.03% for non-model organisms [40].

For 99 newt candidate genes with a known match in another organism, Smith-Waterman alignment of Q3 strings against Swiss-Prot achieved an AUC of 0.94, far exceeding BLAST (0.51) and PSI-BLAST (0.52), and approaching structural alignment’s AUC of 1.0. The perfect AUC score for structure alignment with US-align however is such by definition, as no independent ground truth exists for newt. Therefore homology itself was defined as TM-score above 0.5. The performance of Q3 with an AUC of 0.94 should consequently be read as closely tracking the TM-score-based proxy rather than a validated true homology detection.

The more informative case is the 20 proteins with no existing match at all. Smith-Waterman on Q3 found TM-score-supported hits for 7 of these, outperforming PSI-BLAST on the same query set in 15 of 20 comparisons. For one example the structural match between newt protein mRNA116414 and O50655 suggests the unannotated protein is likely an integrase/recombinase. This is exactly the kind of result Q3 is suited to producing: it prioritizes structurally similar candidates that sequence-alignment-based search misses, generating concrete, testable hypotheses about identity or function. It is not, on its own, evidence of homology or a functional annotation: the newt example is illustrative, producing a promising candidate worth further investigation and not a validated result.

### 4.6 Scope and limitations of the evidence

Several limitations need to be considered before looking to generalize these results.

#### Global AUC versus early retrieval

As established in subsection 4.3, AUC and AUROC1 answer different questions, and a method can score well on one while lagging on the other. Q3’s own AUC vs. AUROC1 divergence across datasets (Table 2) is itself an instance of this.

#### Predicted Q3 tested on one benchmark

As noted in subsection 4.4, predicted Q3 results are reported only for CATH S20; the corresponding CATH and SCOPe40 entries are absent from our benchmark (Table 2) and would be needed before generalizing the finding.

#### Uncertain database labels

Superfamily assignment is a complex process: we found cases where visually near-identical structures were placed in different SCOPe superfamilies (Fig. 2), likely reflecting functional distinctions, the lack of compelling evidence for common ancestry or in rare cases possibly even curation inconsistencies rather than genuine structural dissimilarity. Because our benchmark treats superfamily co-membership as ground truth, such labeling noise places a ceiling on every method’s apparent performance, including US-align’s.

#### Absence of runtime measurements

We report accuracy metrics throughout but did not benchmark runtime or throughput for any method. Claims about the practical efficiency of secondary-structure-based search relative to tertiary methods are based on SOTA literature, e.g. the benchmark reported by van Kempen et al. [11] as well as the hypothesis that applying existing algorithms and systems to the Q3 structural alphabet should come at no substantial extra computational cost. These are therefore not yet empirically supported and should be considered for future work.

#### TM-score-derived newt proxy

As discussed in subsection 4.5, the newt case study lacks independent ground truth and substitutes a TM-score *>* 0.5 threshold for true homology, following precedent [35, 41]. Results on this dataset should be read as agreement with a structural proxy for homology, not as validated homology annotations. Relatedly, our Bayesian-derived Q3 score threshold (0.85) is a heuristic from this same limited setting; deploying it operationally would require calibration on a larger, independently labeled dataset.

### 4.7 Future directions for complementary protein representations

Our results point toward Q3 as one representation among several to be combined and tested systematically, rather than as a replacement for existing tools. Two results from this study illustrate why this needs to be done carefully rather than assumed. First, when we extended Q3 to a 6-letter alphabet by injecting long-range residue contacts to move it closer to 3Di, performance did not improve (AUC 0.90 on SCOPe40, matching basic Q3), so added structural detail does not automatically translate into added discriminative signal, echoing the redundancy argument from subsection 4.2. Second, improving the alignment algorithm itself, that is moving from Levenshtein to Smith-Waterman with BLOSUM-style gap penalties and a purpose-built substitution matrix raised CATH S20 AUC to 0.85, within 5% of the structural-alignment gold standard, showing that scoring refinements can meaningfully close the gap without changing the underlying representation at all.

These two results together motivate a research direction focused on controlled tests of complementarity: systematically combining Q3 with amino-acid sequence, 3Di, 3D coordinates, and protein embeddings, and asking specifically where each contributes independent signal rather than redundant coverage of the same structural information. Beyond representation combination, our Bayesian threshold estimate suggests calibration is a tractable next step toward operational deployment, and our substitution-matrix result suggests scoring refinement is a second lever largely independent of representation choice. Further optimizations such as indexing the database and adding a significance metric, e.g. *E-value*, as demonstrated by search tools such as Foldseek demonstrate further room for improvement.

A recent precedent for this kind of representation-combining approach is Foldseek itself, allowing for usage of both 3Di and AA sequence during search [53], as well as FoldMason [54], which combines primary and tertiary structure for faster multiple structure alignment. This can serve as a proof of concept that representation-combination strategies like the ones proposed here are already worthwhile practical tools.

On the CATH dataset, secondary structure was only 4% behind the gold standard TM-score in AUC, and this gap increased modestly to 9% for the most challenging benchmarks. This performance is particularly noteworthy given that HMM-based tools like HHblits are 2-15% behind tertiary structure alignment accuracy in similar setups. Looking at the score distributions (see S3 Note), it becomes evident that this gap is explained by HHblits’ assigning half of the positives as false negatives close to 0. This is also visible from our example in Fig. 1. Even while aligning the generated HMM sequence profiles, HHblits is virtually unable to find any sequence similarity, while it is visible that the structures are very similar and the secondary structure alignment nicely reveals the common motif.

Now, while we mostly used experimental protein structures to determine secondary structure in our benchmark on protein domains, it is important to note that for any potential application moving forward, one would rather deal with protein sequences that do not have a structure readily available.

Secondary structure assignment itself is subject to variability, with accuracy limits of approximately 88–92% due to so-called “chameleons”—regions of up to 11 residues that can adopt different secondary structure states. Even experimentally determined structures exhibit differences of 5–15% between X-ray and NMR models. [51]. Nevertheless, recent advancements in prediction methods have approached these natural limits, offering robust and reliable secondary structure assignments [55]. The state-of-the-art methods achieve an accuracy of 81 - 89% [20, 38, 52], a performance that, according to our results, seems adequate for constructing meaningful secondary-structure alignments, where interpretability and structural insight can complement sequence-based similarity measures. Our results show, that predicted secondary structure carries virtually the same retention of fold-recognition signal (see Table 2).

Our results demonstrate that secondary structure carries substantial structural information, consistent with earlier studies [12]. This suggests that there is potential for developing hybrid methods that incorporate secondary structure in fields of protein search, protein prediction, functional annotation, domain annotation, and more. But this need not be limited to secondary structure alone.

Evolutionary information has been critical for sequence-based methods like HMMs, but there is untapped potential in integrating evolutionary data into tertiary structure representations. By combining sequence evolution and structural evolution, more sophisticated models could be built to predict structure-function relationships across species.

Our analysis also sheds light on the inherent correlations between protein representations. Secondary structure elements, such as helices and strands, strongly correlate with specific 3Di alphabet characters (e.g., *{V, L, S, C}*_3_*_Di_* for helices). This suggests that information encoded in tertiary representations is often implicitly captured by secondary structure, further validating its utility for RHD tasks.

Emerging representation learning approaches, such as autoencoders and transformers, could further benefit from incorporating secondary structure as an intermediate feature. Tools like Foldseek, which implicitly leverage secondary structure information, already hint at the potential of such comprehensive frameworks.

This could mean that as more high-resolution structural data becomes available, there’s an opportunity to create universal protein representation frameworks. These would integrate all three structural levels into a unified model, making it possible to predict function, interaction, and dynamics from minimal data inputs. An obvious step towards this would be to include the development of multi-level scoring functions that optimise across all structural levels for diverse tasks such as docking, folding, and protein design.

As demonstrated before, sequence-based alternatives such as Foldseek, MMSeqs2, Reseek, BLAST etc. [11, 21, 49, 56] are computationally much more efficient than structure-based methods like TM-align or Dali. Given the sequence nature of the secondary structure strings, it is reasonable to assume, that similar computational performance is possible via algorithmic refinement. In practical scenarios, secondary structure can then serve as a computationally efficient alternative to tertiary structure. For newly sequenced genomes, such as the Iberian ribbed newt in our case study, tertiary structure predictions may be infeasible due to computational costs, and given that the bottleneck is MSA generation, the same holds for HMM-based methods. Secondary structure prediction, on the other hand, is substantially faster [20] and can still enhance RHD performance, as demonstrated by our results. This makes secondary structure a promising basis for functional annotation in large-scale genomic studies.

## 5 Conclusion

Remote homology detection and functional annotation present complex challenges, with the choice of protein representation being crucial. Historically, primary structure has dominated due to its abundance and effective algorithms, but it lacks the spatial insights provided by tertiary structures, which have been limited by data scarcity and computational costs. The advent of tools like AlphaFold, which offers large-scale tertiary structures, and Foldseek, which enables rapid structural searches, has transformed the landscape of protein analysis.

At present, rapid and accurate remote homology detection is possible through the utilisation of tertiary structure, rather than the exclusive reliance on sequences [11]. Currently, accurate remote homology detection is achievable through tertiary structure and the use of embeddings, but the full potential of secondary structure remains underexplored. Our study demonstrates that secondary structure can yield results comparable to full 3D structural alignments in distinguishing superfamily membership.

Moreover, secondary structure offers a unique advantage in flexibility; while tertiary structural alignment algorithms may struggle with conformational variations, secondary structure may be less sensitive to conformational variation than coordinate-based alignment, although we did not test this here. This makes it a promising component for hybrid models, enhancing protein search, classification, and functional prediction. The success of Foldseek may partly stem from its implicit use of secondary structure. Other attempts at such hybrid models have already been made, e.g., FoldMason combining primary structure and tertiary structure representations for faster multiple structure alignments [54].

Future efforts should focus on integrating primary, secondary, and tertiary levels into unified frameworks, leveraging the strengths of each representation to drive robust, scalable protein analysis across diverse applications. In light of these findings, secondary structure warrants renewed attention as a valuable intermediate in computational biology, potentially serving as a cornerstone for integrated methodologies.

## Supporting information

**S1 Note. Custom substitution matrix for Q3 secondary structure.** Pairwise substitution values used in the Smith-Waterman alignment algorithm for SS strings derived using log-odds ratios from Pfam-A Seed alignments.

**S2 Table. Results of searching with SS strings and local alignment.** The optimal TM-scores for PSI-BLAST and SW for all 20 proteins of interest. The reported TM-scores correspond to the highest pairwise TM-score among the 5 highest-scoring matches found with that method for each query. Marked in red are the queries, for which no hits were found with PSI-BLAST and the top hit for SW exceeded TM-score of 0.5.

**S2 Fig. Best matches for 20 queries from Newt.** Each of the 20 unannotated query proteins from newt side by side with their best matching Swiss-Prot structure according to secondary structure SW score. While some pairs show either strong structural similarity or structural similarity between parts of the structure, we also observe cases where the highest-scoring match shares very little structural similarity.

**S3 Note. Score distributions of methods.** Score distributions of all methods; continuous line shows distributions for pairs in the same superfamily, dashed line for those that are in different superfamilies. This allows us to see how well methods separate positive pairs from negative pairs.

**S4 Note. Frequency of secondary structure assignments in CATH and SCOPe40.** The frequency of each letter in the Q8 secondary structure representation according to the DSSP method as well as distribution of domain lengths in the CATH dataset.

**S5 Note. Further analysis of structural alphabets.** The likelihood of replacement for Q3 secondary structure letters (S, H, L) in CATH and SCOPe40, as well as the correlation between protein representation alphabets. Shows heatmap of correlation between Q3, Q8, 3Di alphabet and primary structure representation.

**S6 Note. Further Statistical Analysis.** We calculate the conditional probability for a given secondary structure alignment score (SS score) of protein pairs being in the same superfamily.

