## Supplementary Information for "How informative are structural alphabets for representing protein folds?"

### Supplementary Material

September 29, 2026

#### 1 Supplementary Note S1

##### 1.1 Custom substitution matrix for Q3 secondary structure derived using log-odds ratios from Pfam-A Seed alignments

**Table 1.** Pairwise substitution values used in the Smith-Waterman alignment algorithm for SS strings.

| | H | L | S |
|---|---|---|---|
| H | 2 | | |
| L | -7 | 3 | |
| S | -16 | -5 | 4 |

**Table 2.** In order to verify the robustness of the matrix, we recalculated the matrix for smaller sample sizes of **a)** 500, **b)** 1,000, **c)** 5,000 and **d)** 10,000 alignments that were randomly sampled from the full Pfam-A Seed alignment set. As can be seen from a) and b) for smaller sample sizes, there are minor deviations of magnitude up to 2 in a) and 1 in b) respectively; however, for a sample size of 5,000 the substitution matrix values match the default matrix's values, such that it is safe to assume our matrix to be robust. We ran a similar experiment to test robustness against sequence similarity, where matrices were computed using only alignments of maximum pairwise similarity of 20%, 45%, 50%, 62%, 80%, and 90% respectively. All of these matched the original matrix, which is not surprising since domains are considered to have structurally conserved motifs.

| | H | L | S |
|---|---|---|---|
| H | 1 | | |
| L | -7 | 4 | |
| S | -16 | -3 | 4 |

**(a)**

| | H | L | S |
|---|---|---|---|
| H | 2 | | |
| L | -8 | 3 | |
| S | -16 | -5 | 4 |

**(b)**

| | H | L | S |
|---|---|---|---|
| H | 2 | | |
| L | -7 | 3 | |
| S | -16 | -5 | 4 |

**(c)**

| | H | L | S |
|---|---|---|---|
| H | 2 | | |
| L | -7 | 3 | |
| S | -16 | -5 | 4 |

**(d)**

### 2 Supplementary Note S2

#### 2.1 Results of searching with SS strings and local alignment

**Table 3.** The optimal TM-scores for PSI-BLAST and SW for all 20 proteins of interest. The reported TM-scores correspond to the highest pairwise TM-score among the 5 highest-scoring matches found with that method for each query. Marked in red are the queries, for which no hits were found with PSI-BLAST and the top hit for SW exceeded TM-score of 0.5

| Query | PSI-BLAST hits | TM(PSI-BLAST) | TM(SW) |
|---|---|---|---|
| mRNA10971 | 4 | 0.35 | 0.38 |
| mRNA116398 | 29 | 0.81 | 0.82 |
| mRNA116414 | 0 | NaN | 0.54 |
| mRNA116546 | 985 | 0.35 | 0.40 |
| mRNA126051 | 0 | NaN | 0.69 |
| mRNA131318 | 655 | 0.50 | 0.37 |
| mRNA131915 | 0 | NaN | 0.36 |
| mRNA134396 | 840 | 0.54 | 0.54 |
| mRNA140328 | 1144 | 0.88 | 0.81 |
| mRNA143547 | 817 | 0.59 | 0.37 |
| mRNA152591 | 0 | NaN | 0.48 |
| mRNA152608 | 0 | NaN | 0.35 |
| mRNA32025 | 935 | 0.62 | 0.36 |
| mRNA46480 | 666 | 0.86 | 0.86 |
| mRNA55990 | 2 | 0.33 | 0.37 |
| mRNA5S991 | 0 | NaN | 0.31 |
| mRNA61078 | 661 | 0.44 | 0.46 |
| mRNA61095 | 263 | 0.94 | 0.95 |
| mRNA65919 | 26 | 0.56 | 0.35 |
| mRNA88359 | 708 | 0.43 | 0.43 |

#### 2.2 Best matches for 20 queries from Newt

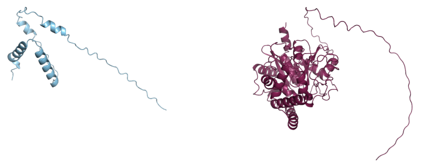

**(a)** Query: mRNA10971, Match: P19922

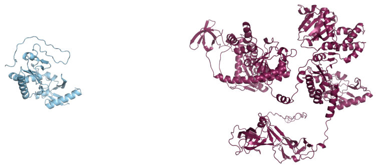

**(b)** Query: mRNA116398, Match: P10394

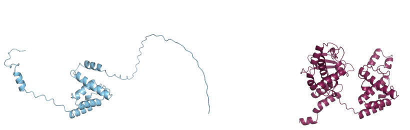

**(c)** Query: mRNA116414, Match: O50655

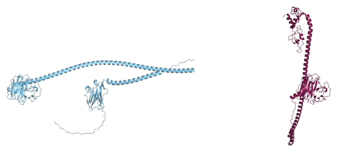

**(d)** Query: mRNA116546, Match: Q6PGR9

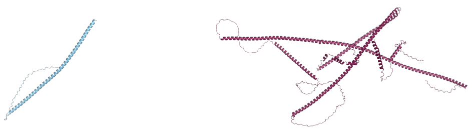

**(e)** Query: mRNA126051, Match: Q95KD7

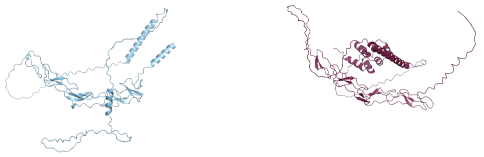

**(f)** Query: mRNA131318, Match: P08138

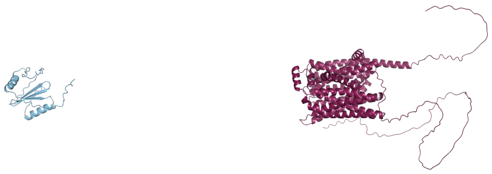

**(g)** Query: mRNA131915, Match: O24381

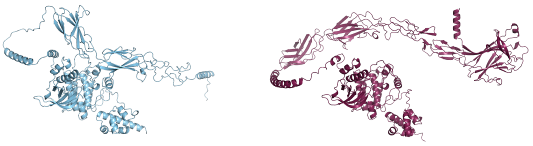

**(h)** Query: mRNA134396, Match: P09759

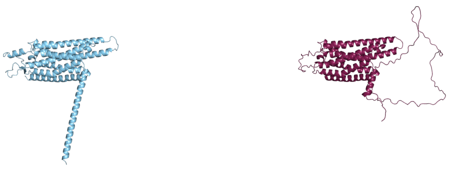

**(i)** Query: mRNA140328, Match: P46616

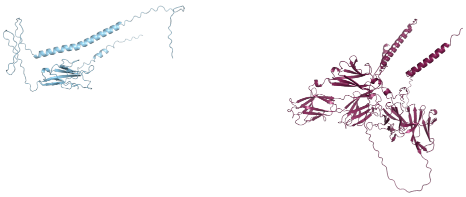

**(j)** Query: mRNA143547, Match: P81265

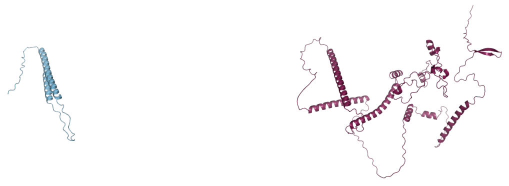

**(k)** Query: mRNA152591, Match: Q9W062

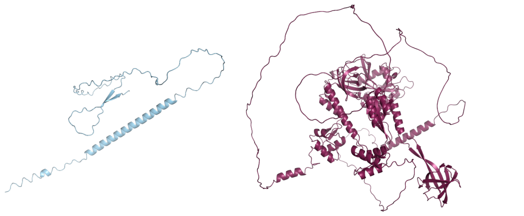

**(l)** Query: mRNA152608, Match: B0RDY9

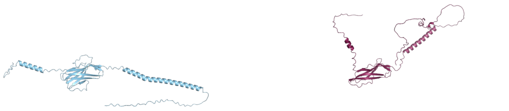

**(m)** Query: mRNA32025, Match: Q8NET5

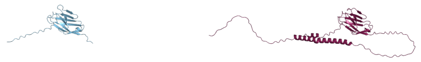

**(n)** Query: mRNA46480, Match: Q6SJQ5

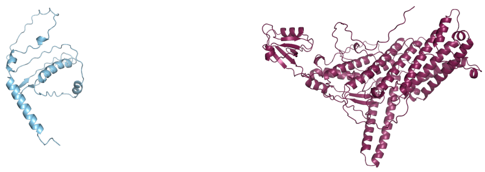

**(o)** Query: mRNA55990, Match: Q8BWR8

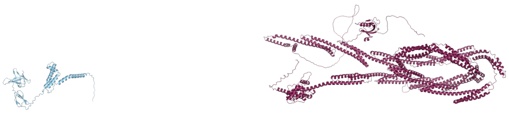

**(p)** Query: mRNA55991, Match: Q62261

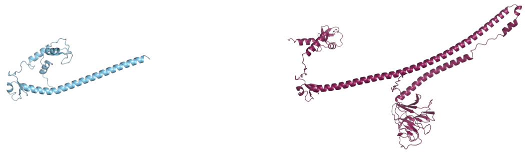

**(q)** Query: mRNA61078, Match: P14373

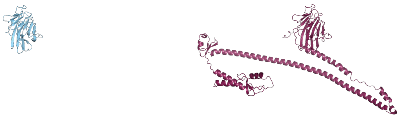

**(r)** Query: mRNA61095, Match: Q5E9G4

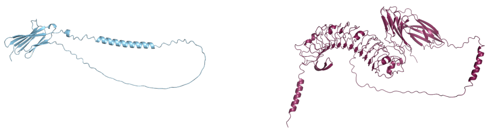

**(s)** Query: mRNA65919, Match: Q9ESY6

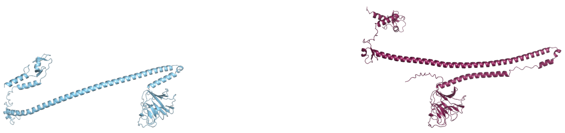

**(t)** Query: mRNA88359, Match: Q62158

**Fig 1.** Each of the 20 unannotated query proteins from newt side by side with their best matching Swiss-Prot structure according to secondary structure SW score. While some pairs show either strong structural similarity or structural similarity between parts of the structure, we also observe cases where the highest-scoring match shares very little structural similarity.

### 3 Supplementary Note S3

#### 3.1 Score distributions of methods

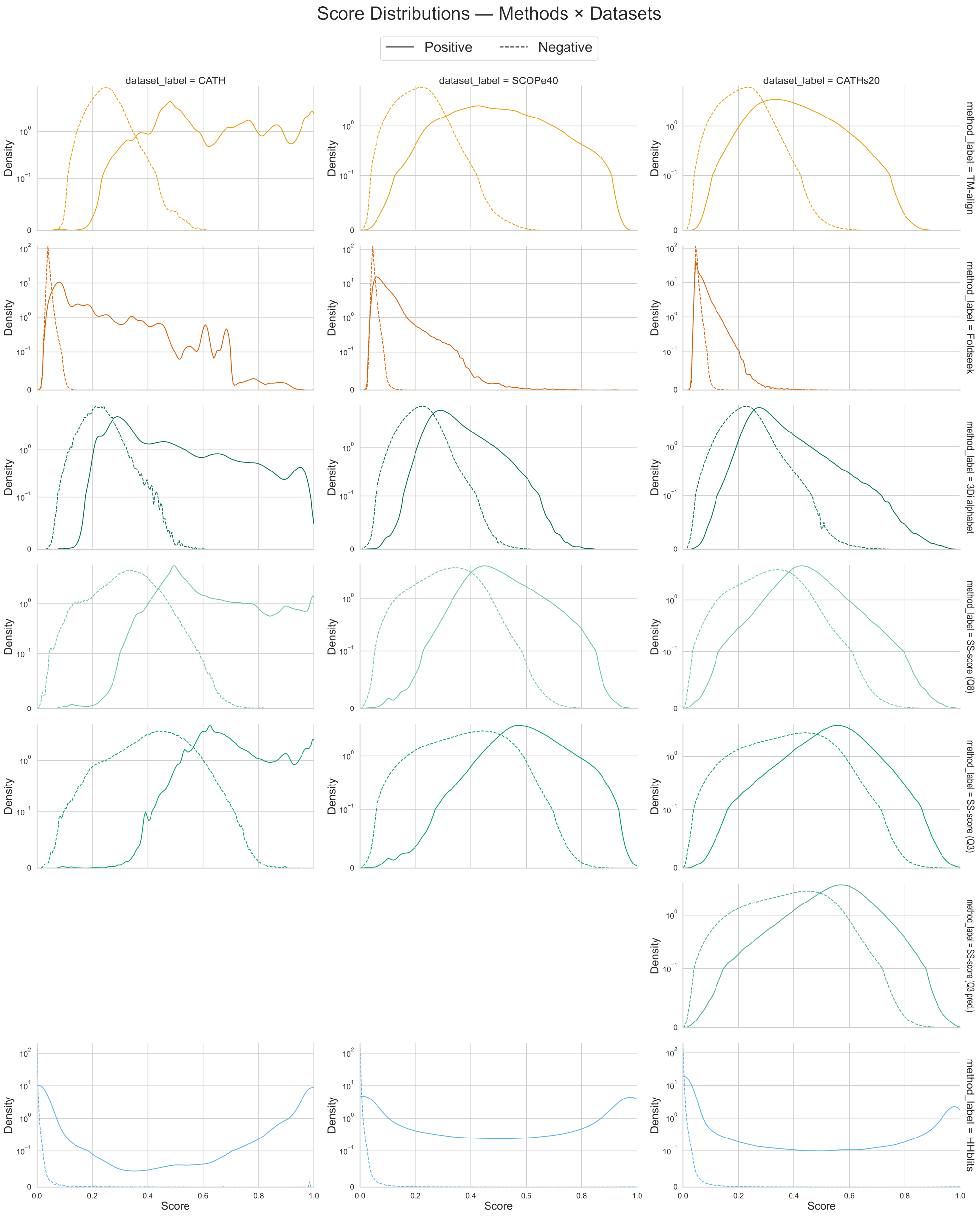


**Fig 2.** Score distributions of all methods; continuous line shows distributions for pairs in the same superfamily, dashed line for those that are in different superfamilies. Note, that distributions have been normalised per group, i.e., for positive pairs and negative pairs each per method and dataset; also, the y-axis is symlog scale, due to dataset imbalance.

This allows us to see how well methods separate positive pairs from negative pairs. Generally we observe the same trends we saw before; difficult datasets are consistently more challenging across all datasets (evident from overlap). What stands out is that while US-align and structure alphabet methods are rather unimodal distributions, HHblits shows a bimodal distribution for the positive pairs, with roughly half of the pairs being scored at 100 % probability of being homologous and the other half at around 0% probability, meaning that in a practical scenario HHblits categorises half of the positives as false negatives, even at the easiest difficulty.

### 4 Supplementary Note S4

#### 4.1 Frequency of secondary structure assignments in CATH and SCOPe40

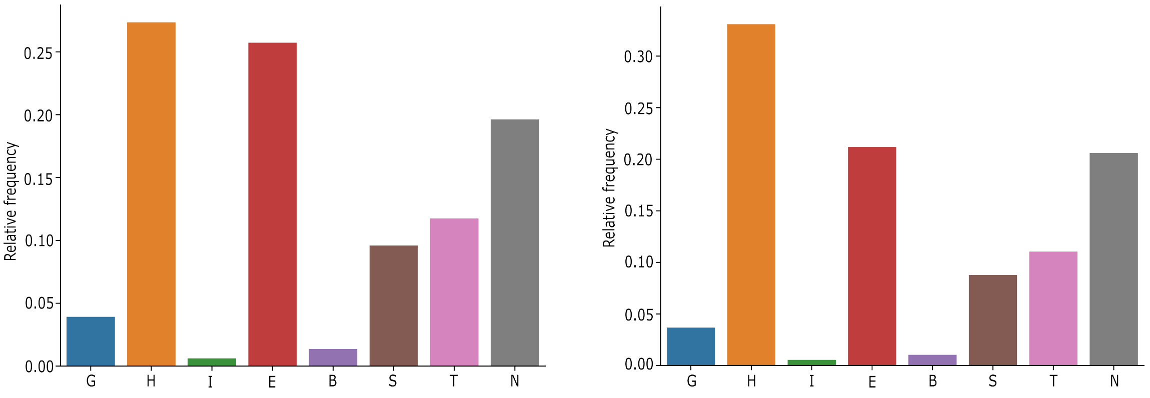


(a) (b)

**Fig 3.** The frequency of each letter in the Q8 secondary structure representation according to the DSSP method. (a) CATH dataset (b) SCOPe40

#### 4.2 Domain length distribution in the CATH dataset

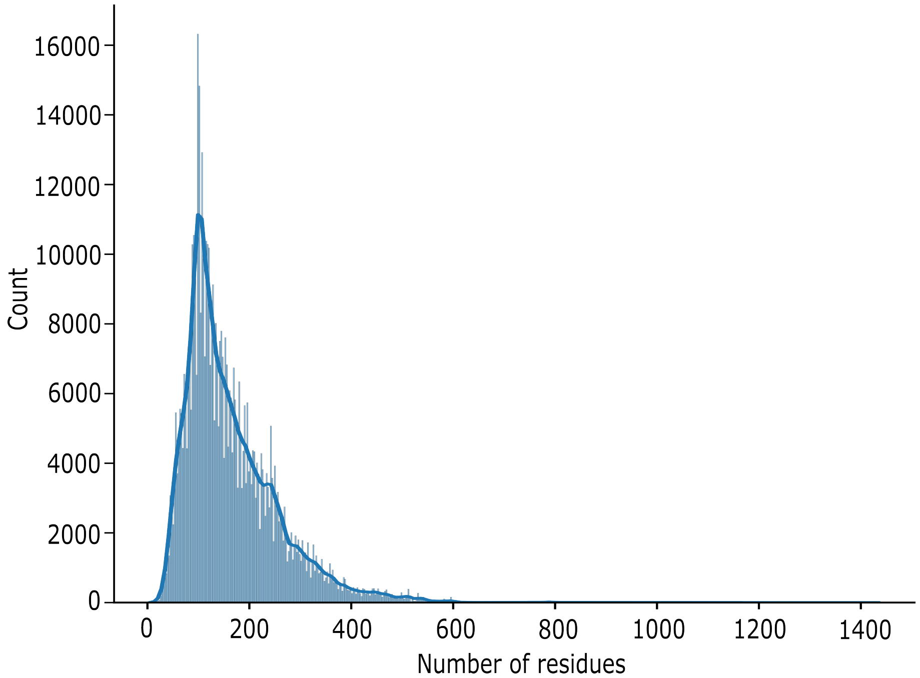


**Fig 4.** Distribution of domain lengths in the CATH dataset. Based on this data, we defined a representative range of amino acids per domain to be between 50 and 250 amino acids.

### 5 Supplementary Note S5

#### 5.1 The likelihood of replacement for Q3 secondary structure letters (S, H, L) in CATH and SCOPe40

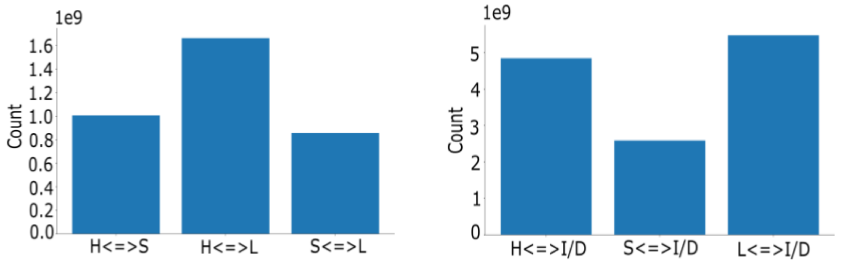


**(A)**

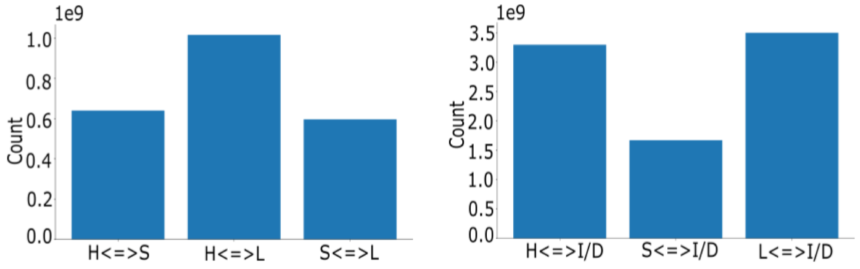


**(B)**

**Fig 5.** Substitution counts for letters in CATH (A) and SCOPe40 (B) datasets. 'H' represents helices, 'S' represents sheets, 'L' represents loops, and 'I/D' stands for insertion/deletion events. The <=> symbol indicates bidirectional substitution. In both panels (A) and (B), the left figures show substitution counts between 'H,'S,' and 'L', while the right figures display substitution counts among 'H,'S,' 'L', and insertions/deletions.

#### 5.2 Correlation between protein representation alphabets

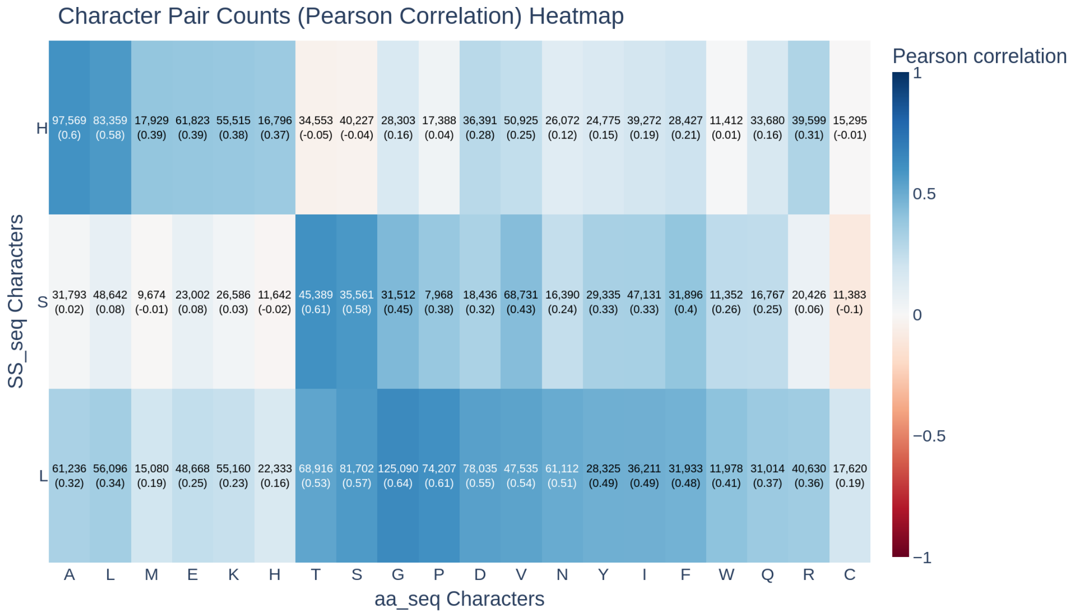


**Fig 6.** Heatmap of correlation between 3-letter secondary structure and primary structure representation. Cells are annotated by total occurrence of this pairing, Pearson correlation in brackets. Colour according to Pearson correlation.

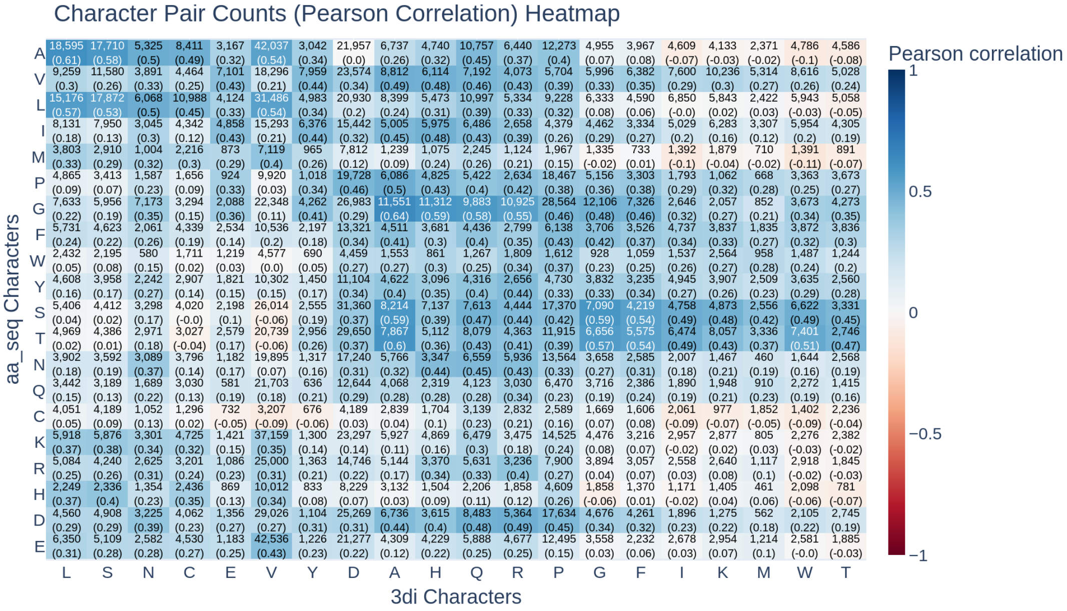


**Fig 7.** Heatmap of correlation between 3Di alphabet and primary structure representation. Cells are annotated by total occurrence of this pairing, Pearson correlation in brackets. Colour according to Pearson correlation.

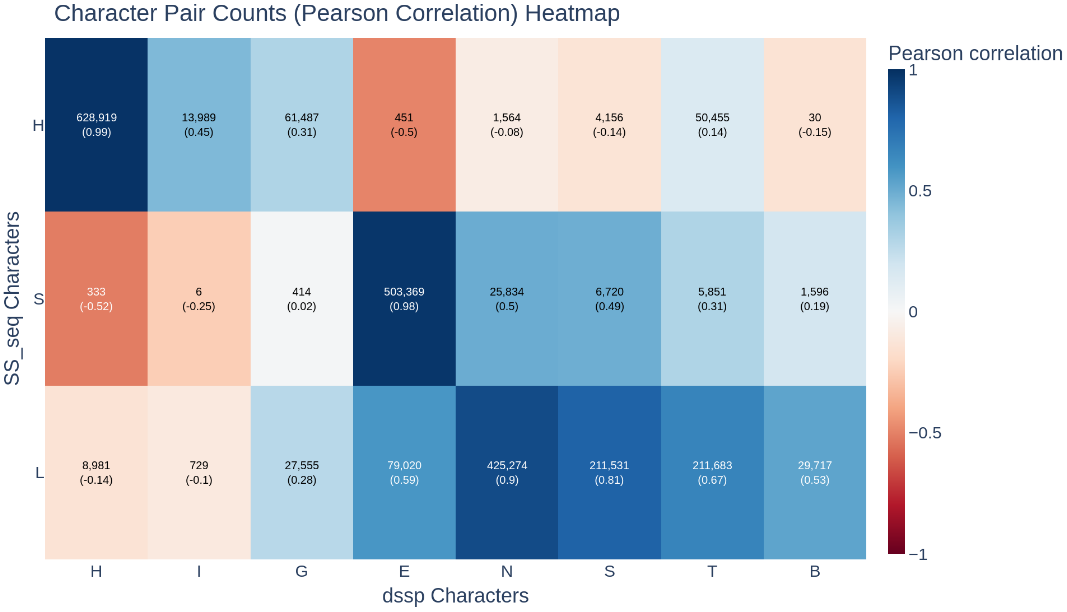


**Fig 8.** Heatmap of correlation between Q3 and Q8 secondary structure representation. Cells are annotated by total occurrence of this pairing, with Pearson correlation in brackets. Colour according to Pearson correlation.

### 6 Supplementary Note S6

#### 6.1 Further Statistical Analysis

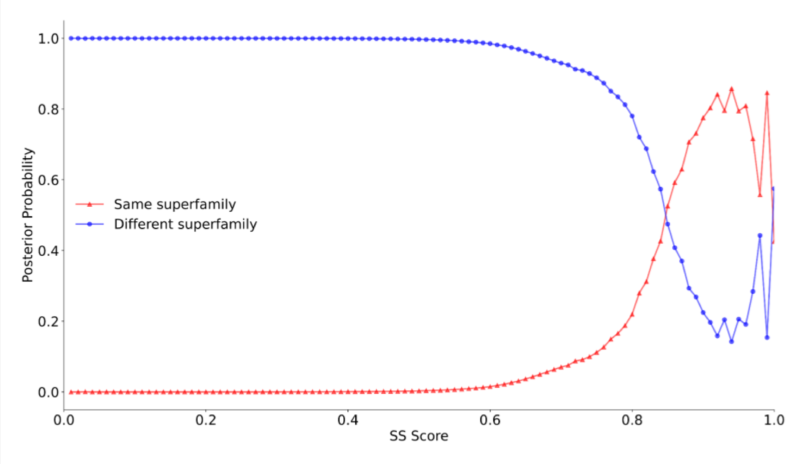


**Fig 9.** The posterior probabilities of SCOPe domain pairs for a given secondary structure score being in the same superfamily (red) or different superfamilies (blue). Both lines cross at around 0.85.

We calculate the conditional probability for a given secondary structure alignment score (SS score) of protein pairs being in the same superfamily. For an SS score below 0.6, the probability of a protein pair belonging to the same superfamily is close to 0; this is the case for only a few pairs. However, we can also observe that for an SS score above 0.8, that probability increases sharply to around 50%. Around the score of 0.85, we observe a clear phase transition. There are also a few unexpected spikes or respective drops, especially for SS scores around 1.0, but these are rather outliers and can be explained by anomalies in and the reduced size of the dataset. Nonetheless, this phase transition points at a possible threshold of 0.85, indicating that two proteins can be expected to be in the same superfamily above that threshold. The same procedure is done on the TM score in our data, and as was underlined in previous studies [1], the threshold of 0.5 is optimal for superfamily classification, as shown in the figure below. We also performed the same analysis on the SCOPe dataset and for several methods, i.e., TM score and SS score, using a Q3 secondary structure assignment, consisting of a three-letter alphabet, and the Q8 secondary structure assignment, consisting of an eight-letter alphabet [2,3].
